# Integrated single-cell and spatial transcriptomic analyses unravel the heterogeneity of the prostate tumor microenvironment

**DOI:** 10.1101/2022.03.18.484781

**Authors:** Taghreed Hirz, Shenglin Mei, Hirak Sarkar, Youmna Kfoury, Shulin Wu, Bronte M. Verhoeven, Alexander O. Subtelny, Dimitar V. Zlatev, Matthew W. Wszolek, Keyan Salari, Evan Murray, Fei Chen, Evan Z. Macosko, Chin-Lee Wu, David T. Scadden, Douglas M. Dahl, Ninib Baryawno, Philip J. Saylor, Peter V. Kharchenko, David B. Sykes

## Abstract

The treatment of primary prostate cancer delicately balances an active surveillance approach for low-risk disease with multimodal treatment including surgery, radiation therapy, and hormonal therapy for high-risk disease. Recurrence and development of metastatic disease remains a clinical problem, without a clear understanding of what drives immune escape and tumor progression. Here, we sought to comprehensively describe the tumor microenvironment of localized prostate cancer contrasting this with adjacent normal samples and healthy controls. We performed single-cell RNA sequencing and high-resolution spatial transcriptomic analysis. This revealed tumor context dependent changes in gene expression. Our data point towards an immune suppressive tumor microenvironment associated with suppressive myeloid populations and exhausted T-cells, in addition to high stromal angiogenic activity. We inferred cell-to-cell relationships at an unprecedented scale for ligand-receptor interactions within undissociated tissue sections. Our work provides a highly detailed and comprehensive resource of the prostate tumor microenvironment as well as tumor-stromal cell interactions.

**Highlights:**

- Characterization of prostate cancer by combined scRNA-seq and spatial transcriptomic analysis
- Primary prostate cancer establishes a suppressive immune microenvironment
- The prostate tumor microenvironment exhibits a high angiogenic gene expression pattern
- A new computational analysis pipeline to deconvolute context-specific differential gene expression

## Introduction

Localized prostate cancer is a clinically heterogeneous disease. Some patients present with indolent low-risk prostate tumors that can safely be observed, while others have aggressive high-risk disease that carries a substantial relapse risk even following state-of-the-art treatment. Despite efforts aimed at early detection and improving our current curative-intent therapies, many patients unfortunately experience recurrence and disease progression (1). There remains a significant need to further our understanding of prostate cancer, where biological insights of the prostate tumor microenvironment (TME) may help to identify novel therapeutic targets. We examined the supportive cellular states and molecular relationships within the prostate TME to identify changes that drive tumor growth.

Single-cell gene expression technologies have made it possible to assess thousands of cells within a single sample, revealing subtleties in tumor cell heterogeneity as well as a complex TME (2–4). Examinations of normal adult human prostate gland (5) and prostate cancer have provided detailed descriptions of the epithelial and tumor cells as well as cell states in both prostate adenocarcinoma (6–9) and neuroendocrine tumors (10). However, the immune cells within the prostate microenvironment have not been rigorously characterized at the single-cell level. The prostate TME typically contains few immune cells, and it is hypothesized that this feature may explain the generally poor response of prostate cancer to immunotherapy (11, 12). We therefore processed fresh prostate and tumor samples using a method that enriched and preserved immune cell populations so to characterize the immune microenvironment at high-resolution.

To validate our single-cell findings, we used a parallel spatial transcriptomic technique (Slide-seqV2), where the tissue architecture and cell-cell proximity relationships are preserved (13, 14). We thus also characterized the spatial organization of tumors from patients with low-risk and high-risk prostate cancer. In addition, we developed a new computational means of data analysis to examine the transcriptional impact of tumor cells on neighboring stromal cells, including fibroblasts, pericytes and endothelial cells.

Together, this work provides a compendium of the prostate TME with a particular focus on immune populations. We further reveal the transcriptional state of stromal cells based on their spatial localization within the tumor. In sum, our data reveal a highly immune suppressive TME and describe tumor-induced alterations of neighboring cells that promote tumorigenesis and progression.

## Results

### The prostate TME characterized by single-cell and spatial transcriptomic analysis

Fresh prostate cancer samples were collected from 19 treatment-naïve patients diagnosed with prostate adenocarcinoma and undergoing radical prostatectomy. In 14 of the 19 patients, matched ‘normal’ benign prostate gland tissue adjacent to the tumor was also sampled. As controls, samples from prostate tissue not harboring cancer were collected from 4 patients (undergoing cystoprostatectomy for bladder cancer), and one healthy prostate was collected as part of a rapid autopsy from a patient with metastatic non-small cell lung cancer (**Figure 1A**).

**Figure 1.**
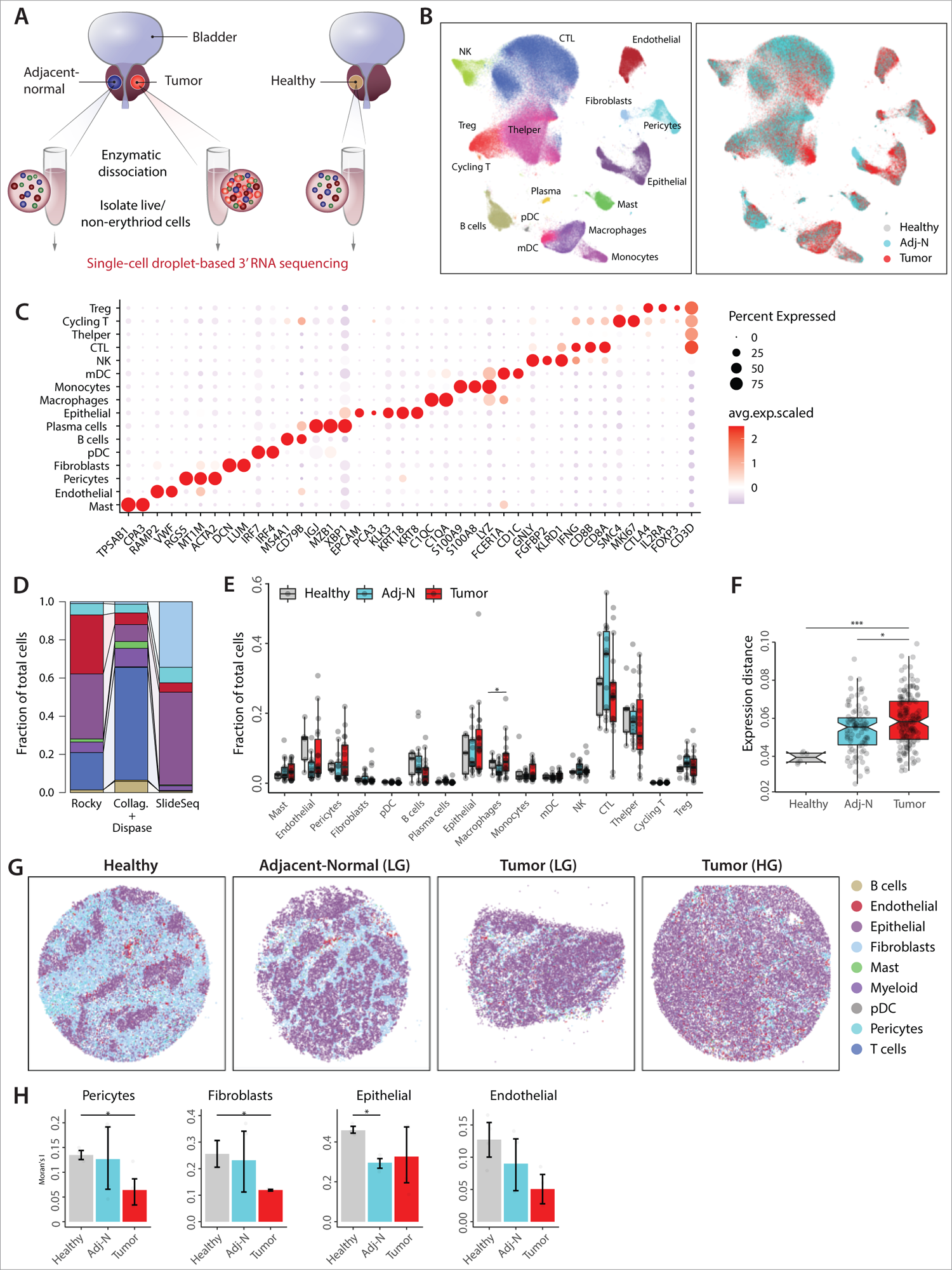
The prostate TME characterized by single-cell and spatial transcriptomic analysis. **A.** Schematic illustration of sample collection and processing. **B.** Integrative analysis of scRNA-seq samples visualized using a common UMAP embedding for cell annotation (left) and sample fractions (right). **C.** Dotplot representing key-marker gene expression in major cell types. The color represents scaled average expression of marker genes in each cell type, and the size indicates the proportion of cells expressing marker genes. **D.** Stacked barplots showing the fractional composition of cell number for different clusters within scRNA-seq (using two different dissociation protocols: Collagenases+Dispase and Rocky, see text) and Slide-seqV2. The connection between the stacked barplots connects same cell clusters. **E.** Boxplot comparing proportion of major cell populations between healthy prostate tissues and tissues collected from cancerous prostates (tumor and adj-normal). Significance was assessed using Wilcoxon rank sum test (*p<0.05). **F**. Boxplot showing inter-individual gene expression distances (based on Pearson correlation) within healthy, adj-normal, and tumor samples, averaged across all cell types. Significance was assessed using Wilcoxon rank sum test *p<0.05, ***p<0.001). Boxplots in (E-F) include centerline, median; box limits, upper and lower quartiles; and whiskers are highest and lowest values no greater than 1.5x interquartile range. **G**. Spatial presentation at a high-resolution level using Slide-seqV2 for the major cell populations in healthy, adj-normal of LG case and two tumor tissues collected from a low-grade (Tumor-LG) and high-grade (Tumor-HG) patients. **H.** Barplots showing spatial autocorrelation (Moran’s I) of fibroblasts and pericytes in Healthy, adj-Nomral and Tumor samples. Moran’s I evaluates whether the cells are clustered (high Moran’s I score) or dispersed (low Moran’s I score). Statistical analysis was performed using Wilcoxon rank sum test. (*p<0.05, error bars: SEM).

The cellular composition of the prostate TME was examined across a spectrum of primary tumor grades and stages (pathologic T-stage 2a to 3b; Gleason score 6-10). Samples were divided into low-grade (LG, Gleason 6 and 7, 12 cases) and high-grade (HG, Gleason 8-10, 7 cases) (**Table S1**). Live, non-erythroid cells (DAPI^neg^/CD235^neg^) were collected by fluorescence-activated cell sorting (FACS) from healthy prostate tissues (n=5), prostate tumor tissues (n=12 LG and n=7 HG) and adjacent non-tumor involved prostate tissues (n=11 LG and n=4 HG, hereafter ‘adjacent-normal’). From 14 patients we collected paired tumor tissue and adjacent-normal tissue samples (n=10 LG and n=4 HG) (**Table S1**). All patients had standard pathologic evaluation to confirm their diagnosis (**Figure S1A**).

The transcriptomes of 179,359 single cells were analyzed (average of 4,721 cells per sample and 50,416 transcripts per cell, **Table S2**). Conos (15) (Clustering On Network Of Samples) aligned the samples, and the analysis of the resulting joint cell clusters revealed a rich repertoire of immune cells and non-immune stromal cells (**Figure 1B**). Cell types were annotated based on cell type-specific gene markers, forming 16 major clusters (**Figure 1C, S1B, Table S3).**

Of note, our dissociation protocol was optimized to enrich for immune cells. This was an intentional choice to focus on the prostate immune TME with the goal of understanding why prostate cancers are considered poorly immunogenic and so rarely respond to immunotherapy (16). In comparing our tissue processing method (Collagenases+Dispase) to a published protocol of a single-cell prostate study (Rocky) (5), the Collagenases+Dispase released a higher proportion of immune cells (**Figure 1D, S1C**). Reassuringly, cells liberated by both dissociation protocols showed similar transcriptome profiles (**Figure S1D**).

In terms of the abundance of major cell populations, significant but small absolute differences were observed at the global level in plasma cells, macrophages, and endothelial cells when comparing the tumor sample to the adjacent-normal sample **(Figure 1E)**. Stratifying low-grade (LG) and high-grade (HG) cases, there were similar small but significant changes in plasma cells (adj-normal vs tumor, LG), macrophages (adj-normal vs tumor, LG) and endothelial cells (Healthy vs. adj-N LG) (**Figure S1E**). The few significant differences in cell abundance were likely due to high patient-to-patient variability even within patients who had the same Gleason score **(Figure S1F)**.

The overall similarity of the transcriptional state between samples was examined using a weighted expression distance, revealing a significant increase in the inter-patient variability among the tumor fraction, compared to the adj-normal and healthy fractions **(Figure 1F)**. This suggests divergent trajectories of the cellular states in the tumor region among different patients.

To validate single-cell findings with a dissociation-free approach that preserves tissue architecture, we performed spatial transcriptomics using Slide-seqV2 (13, 14). This provided the opportunity to examine tumor organization at high spatial resolution. Fresh-frozen 10-micron sections were sampled from a healthy prostate sample and two prostate tumor samples (one low grade and one high grade) as well as their corresponding adjacent-normal tissues **(Figure 1G)**.

Robust Cell Type Decomposition (RCTD) was used to assign cell type annotations based on scRNA-seq reference data (**see Methods**) (17). Hallmark genes denoting different cell populations were used to verify the RCTD annotation **(Figure S1G)**. As expected, Slide-seqV2 measurements showed more pronounced differences in cell proportions as compared to the scRNA-seq data, with greatly expanded epithelial and fibroblast populations and a significantly smaller fraction of immune cells (**Figure 1D**).

The cellular architecture viewed through the lens of Slide-seqV2 was reassuringly consistent with what one would expect from standard H&E staining. The highly detailed spatial configuration of the healthy prostate tissue demonstrated well-organized prostate epithelial glands surrounded by immune and non-immune stromal cells including fibroblasts, pericytes, mast cells, and endothelial cells **(Figure 1G, panel 1)**. This architecture was notably disrupted in the cancerous prostate (**Figure 1G, panels 3 and 4**). Differences in tissue organization were quantified by spatial autocorrelation using Moran’s I score, which evaluates the extent to which the cells are clustered (high score) or dispersed (low score) (18). The Moran’s I score for fibroblasts, endothelial cells, and pericytes significantly decreased in tumor as compared to healthy tissues (**Figure 1H**).

### A Prostate Tumor Gene Signature distinguishes normal and malignant luminal epithelial cells

Unsupervised clustering revealed four epithelial subpopulations: basal, luminal, club, and hillock (**Figure 2A**) as denoted by key marker gene expression (**Figure S2A**). Hillock and club cells were identified as transitional cells in a cellular atlas of the mouse lung (19). These cells have also been reported in human prostate tissue (5, 20) and in benign human prostate organoids (7), but their role in prostate tumorigenesis remains unclear.

**Figure 2.**
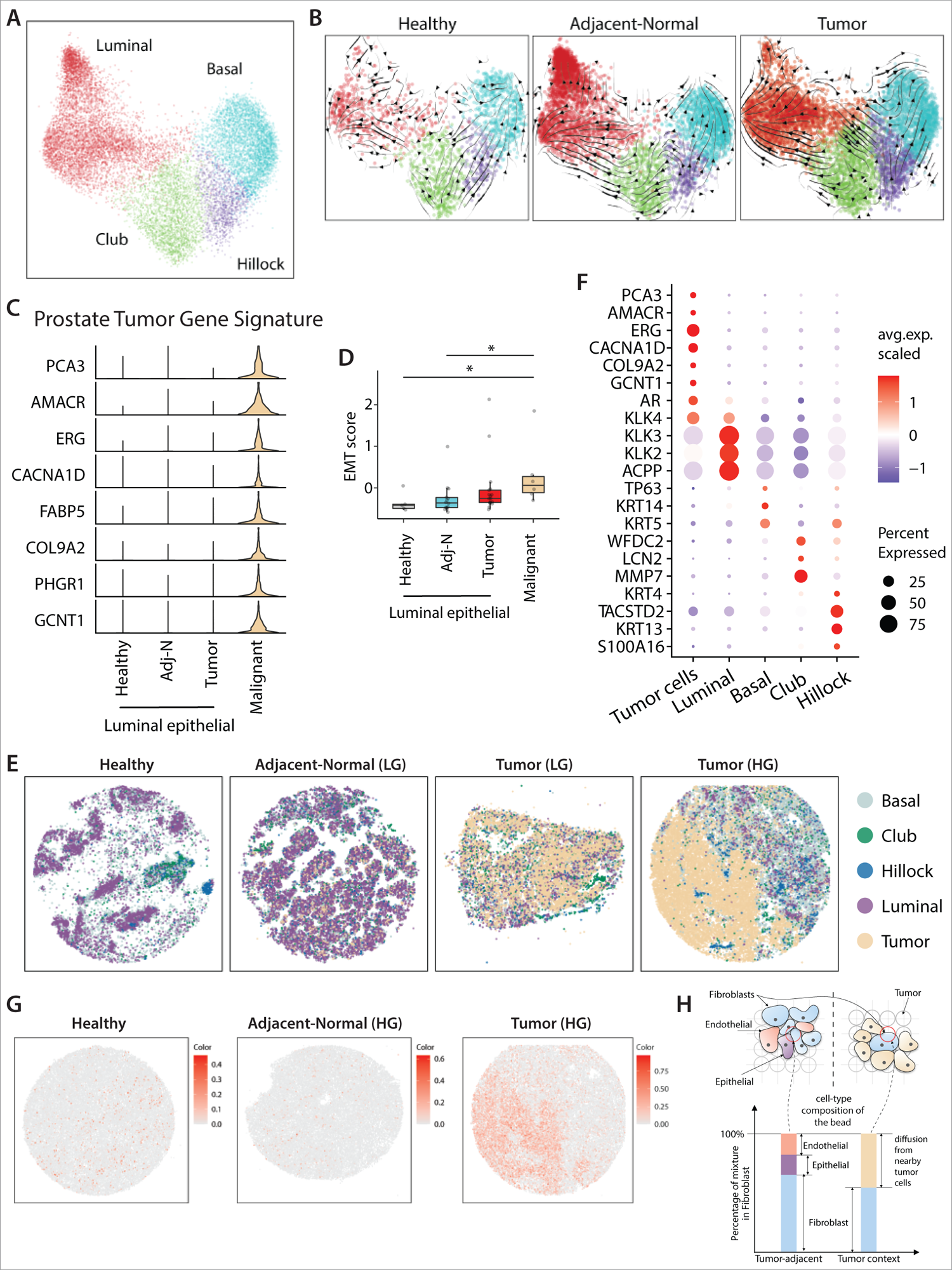
A Prostate Tumor Gene Signature distinguishes normal and malignant luminal epithelial cells. **A.** Joint embedding represent the detailed annotation of epithelial subpopulations in prostate tissues. **B.** RNA velocity analysis of the transitions of epithelial cells, estimated on different sample fraction. **C.** Violin plot showing the expression of genes panel of “Prostate Tumor Gene Signature” in malignant cells and in the epithelial luminal cells of healthy, adj-normal and tumor prostate samples. **D.** Boxplot representing the epithelial-mesenchymal transition (EMT) score in malignant cells and the epithelial luminal cells of healthy, adj-normal and tumor prostate samples. Significance was assessed using Wilcoxon rank sum test (*p<0.05). **E.** Spatial presentation of epithelial subpopulations in healthy, adj-normal (Adj-Normal LG) and two tumor tissues collected from a low-grade (Tumor (LG)) and high-grade (Tumor (HG)) patients. **F.** Dotplot representing key-marker gene expression in epithelial subpopulations in Slide-seqV2. The color represents scaled average expression of marker genes in each cell type, and the size indicates the proportion of cells expressing marker genes. **G**. Spatial presentation for “Prostate Tumor Gene Signature” average expression in healthy, adjacent-normal (HG) and tumor (HG) Slide-SeqV2 pucks. **H.** A schematic view the admixture problem in the Slide-seqV2 puck. The barplot shows the cell-type composition in two different contexts within the same puck. The barplot related to the tumor context contains substantial admixture from nearby tumor cells whereas the one related to tumor-adjacent context is a heterogeneous mixture of different cell-types.

We used RNA velocity to infer the likely trajectories of epithelial cell differentiation (21, 22). One trajectory suggested that club cells act as luminal cell progenitors, an observation previously reported in prostate cancer (23). A second distinct trajectory showed consistent directional flow suggesting that hillock cells may be acting as progenitors for basal cells (**Figure 2B**). Differential gene expression comparing healthy and tumor-associated hillock and club cells showed enrichment in genes involved in urogenital system development and epithelial tubes morphogenesis, respectively (**Figure S2B**) and these cells are known to be enriched in urethra and peri-urethral prostate zones (5).

Malignant cells did not cluster separately from the non-malignant epithelial populations from which they originated. To distinguish malignant cells from normal epithelial cells within the prostate tumor samples, we applied inferCNV (3,24,25) on the four epithelial subpopulations, taking their corresponding subpopulation from healthy samples as a reference. Only cells within the luminal subpopulation showed clear chromosomal aberrations, indicating that the malignant cells are of luminal origin, consistent with previous studies (26) (**Figure S2C**).

Chromosomal aberrations and inferCNV analysis allowed us to separate malignant luminal cells (with genomic aberrations) from normal luminal cells within the tumor. DEG analysis was used to identify an expression signature for the malignant cells, leading to a signature composed of eight genes, which we termed the “Prostate Tumor Gene Signature” **(Figure 2C)**. We applied this gene signature to published bulk RNA-seq of prostate tissues, demonstrating a consistent ability to distinguish tumor samples from adjacent normal samples across four independent datasets (**Figure S2D**) (27–30).

Since we were able to distinguish malignant cells from normal epithelial cells within tumor samples, we assessed for heterogeneity. Independent component analysis (ICA) of malignant cells revealed three major aspects of malignant clusters **(Figure S3A).** Gene Ontology (GO) pathway analysis showed an enrichment in cell growth and epithelial cell migration related genes in malignant cluster 1 (C1) **(Figure S3B)**. Cluster 1 also showed high expression of EGR1, IER2 and KLF6 genes **(Figure S3A)** suggesting roles in prostate cancer progression, motility, and metastasis (31, 32).

Epithelial-mesenchymal transition (EMT) plays an important role in prostate cancer progression and metastasis (33). Malignant cells showed significantly higher EMT gene signature (34, 35) (**Table S4**) as compared to non-malignant luminal cells from the three different sample types (healthy, adj-normal and tumor) (**Figure 2D, Figure S3C**).

Spatially, the healthy prostate demonstrated an organized glandular epithelium with a well-structured bilayer of basal and luminal cells **(Figure 2E).** The adj-normal sample differed with an expansion of the luminal epithelial population, and loss of the well-organized glands (**Figure S1A, Figure 2E**). Epithelial subpopulations were annotated using RCTD and validated using epithelial cell-type specific marker genes (**Figure 2E and F**). The normal clusters of club and hillock cells were disrupted in the tumor and adj-normal samples as demonstrated by spatial autocorrelation (**Figure S3D**).

The “Prostate Tumor Gene Signature” obtained from the single cell experiments was applied to the Slide-seqV2 results. This eight-gene tumor signature successfully identified tumor cells collected from the HG case (**Figure 2G**). Almost no such cells were annotated in the healthy and adj-normal samples (**Figure 2G**), speaking to the accuracy of this “Prostate Tumor Gene Signature”.

### Context-dependent differential expression with linear admixture correction

The edge, or boundary, of the expanding tumor was particularly evident in the HG sample, which could be segmented into two distinct spatial contexts. The tumor context was dominated by dense accumulation of tumor cells, while the tumor-adjacent context was composed primarily of non-malignant epithelial cells (**Figure S3E**). The small fraction of tumor cells detected within the adj-normal sample likely represents real infiltration of tumor cells. Slide-seqV2 allows one to examine the differences in cellular state associated with precise spatial contexts. Annotation tools such as RCTD (17) estimate the fractions of cell types contributing to each bead and identify relatively pure beads that can be confidently assigned to a specific cell type. However, even “pure” beads can carry admixture of transcriptomes from the neighboring cells (**Figure 2H**).

As composition of the cellular neighborhoods varies between different tissue contexts, such admixture will heavily bias transcriptional comparisons of cellular state between contexts. To overcome this admixture effect, we developed a new computational approach which regressed out context-dependent differences that could be attributed to admixture from other cell types, focusing on the residual differences that likely reflect the context-dependent change in the transcriptional state of the target cell-type (**Supplementary Note 1**). In subsequent sections, we apply this approach to contrast the state of the stromal populations between tumor and tumor-adjacent contexts.

### The prostate tumor microenvironment exhibits high endothelial angiogenic activity

The non-immune stroma includes fibroblasts, endothelial cells and pericytes, representing important components of the TME whose function and abundance varies significantly between cancer types (36). We identified five stromal subpopulations including two endothelial, two pericyte, and one fibroblast subpopulation **(Figure 3A)** annotated based on key marker gene expression (37–41) **(Figure S4A and Table S3)**.

**Figure 3.**
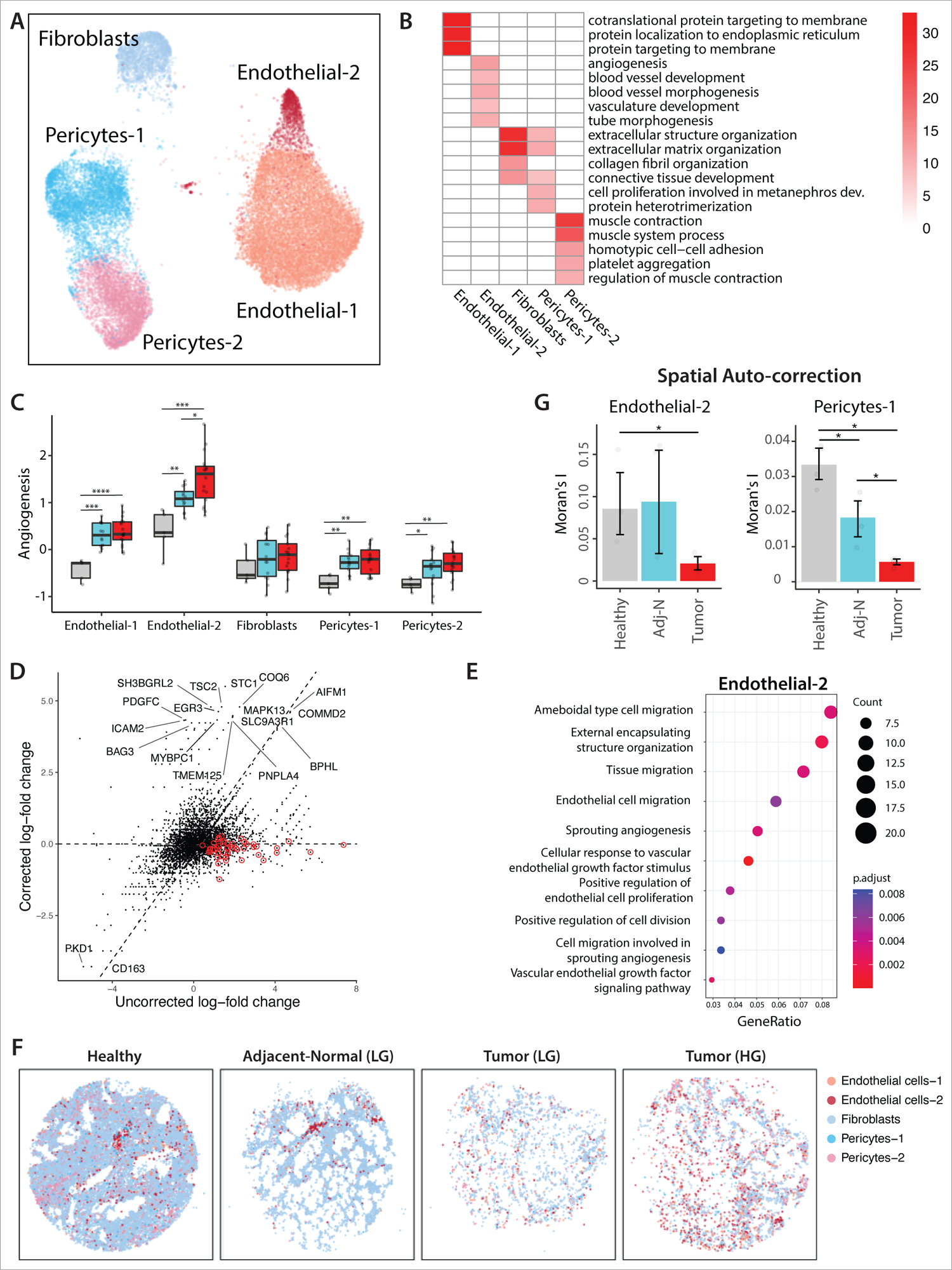
The prostate tumor microenvironment exhibits high endothelial angiogenic activity. **A.** Joint embedding represent the detailed annotation of stromal cells in prostate tissues. **B.** Overview of enriched GO terms of top 200 upregulated genes for each stroma subpopulation based on single-cell data analysis. **C.** Boxplot comparing the angiogenesis signature across the three different sample fractions for each stroma subpopulation. See Supplementary Table S4 for the genes defining angiogenesis signature. Boxplots in include centerline, median; box limits, upper and lower quartiles; and whiskers are highest and lowest values no greater than 1.5x interquartile range. Statistical significance was accessed using Wilcoxon rank sum test (*p<0.05, ****p<0.0001). **D.** The scatterplot showing the effect of linear model-based correction on Endothelial-2 cells. Red dots indicate tumor marker genes. The x-axis is the log-fold change of the genes without the correction, the y-axis is the same after the correction. The top DE genes are text-labeled. **E.** Dotplot shows enriched GO terms of upregulated genes in Endothelial-2 cells in tumor context compared to tumor-adjacent context. **F.** Spatial presentation at a high-resolution level using Slide-seqV2 for the stroma subpopulation healthy, adj-normal (Adj-Normal LG) and two tumor tissues collected from a low-grade (Tumor (LG)) and high-grade (Tumor (HG)) patients. **G.** Comparison of spatial autocorrelation (Moran’s I) of Endothelial-2 cells and Pericytes-1 cells in healthy, adj-normal and tumor samples.

Endothelial-1 cells showed high expression of SELE/SELP/CLU/PLVAP, characteristic of sinusoidal endothelial cells whereas Endothelial-2 cells expressed common arterial genes (HEY1/IGFBP3/FBLN5) (42–46) **(Figure S4A)**. Gene Ontology (GO) analysis of Endothelial-2 cells pointed to pathways involved in blood vessel development and angiogenesis **(Figure 3B).** An angiogenesis gene signature (35) **(Table S4**), demonstrated that the tumor-associated Endothelial-2 cells scored highest when compared to the other stroma populations and when comparing healthy and tumor across almost all populations (**Figure 3C).** The angiogenesis scores of different stromal subpopulations did not differ between LG and HG tumor samples.

Transcriptomic changes of the Endothelial-2 cells were examined within the Slide-seqV2 spatial transcriptomic platform (**Figure 3D**) comparing the ‘tumor’ and ‘tumor-adjacent’ contexts (**Figure S3B**), Pathway enrichment analysis was consistent with the single-cell data of the tumor, showing upregulation of sprouting angiogenesis and vascular endothelial growth factor pathways (**Figure 3E and S4C**).

Endothelial-2 cells in the tumor context also showed upregulation of cell migration and proliferation pathways. This is consistent with the dispersed organization of the Endothelial-2 cells within the tumor tissue in contrast to well-organized structures of the adj-normal and healthy samples (**Figure 3F),** and this was quantified by spatial autocorrelation analysis **(Figure 3G**). Overall, this highlights the relevance of endothelial cells to tumor vascularization and migration, which correlates with prostate cancer disease progression (47).

Perivascular pericytes are another component of the vascular system. These cells exhibit mesenchymal features with multipotency (48), and their role in vasculature development is established while their role in cancer progression is unclear. We identified two pericyte subpopulations **(Figure 3A).** The expression pattern in Pericyte-1 cells was enriched for pathways involved in extracellular structure organization and connective tissue development, while Pericyte-2 cells demonstrated gene signatures enriched for muscle contraction consistent with vascular smooth muscle cells (VSMCs) (**Figure 3B**). In addition, there was a significant increase in the angiogenic gene signature of both pericyte subpopulations in samples collected from cancerous prostate as compared to healthy prostate **(Figure 3C)**. Spatially, Pericyte-1 cells were dispersed in the tumor samples when compared to healthy and adj-normal samples (**Figure 3F and 3G)**. Taken together, these data suggest a role for pericytes in angiogenesis and in remodeling the tumor stroma during prostate cancer progression.

Cancer-associated fibroblasts (CAFs) play a critical role in shaping the TME by promoting tumor proliferation and metastasis (49), enhancing angiogenesis (50), and mediating immunosuppression (51). CAFs are associated with poor prognosis in many cancer types (52–54). In prostate cancer, CAFs play a causal role in cancer development at early disease stages, contributing to therapy resistance and to metastatic progression (55). Fibroblast gene expression patterns showed an enrichment for collagen fibril organization, extracellular structure organization and connective tissue development pathways **(Figure 3B**). These same pathways were also identified within the Slide-seq differential gene analysis, comparing the tumor to the tumor-adjacent context (**Figure S4D**). These data suggest a role for fibroblasts in inducing extracellular matrix remodeling in prostate TME, which in turn is important for tumor progression.

### Coordination between tumor cells and stromal compartment in tumor context

We utilized Slide-seqV2 spatial information to examine potential channels of communication between cells within the tumor ecosystem. While the importance of cell-to-cell signaling is appreciated, it is challenging to infer which cells communicate with each other and via which channels (56). Prediction of possible relationships is based on the expression of ligand and cognate receptor pairs and typically results in many potential interactions; additional filters are needed to distinguish functionally relevant channels. We reasoned that spatial proximity might be one such filter to identify relevant interactions.

We asked whether the corresponding ligand and receptor genes exhibited cooperative upregulation in cells positioned directly next to each. Slide-seqV2 data was used to graph physically adjacent cells, which permitted testing whether a ligand-receptor (LR) score, defined as a product of the two corresponding expression levels, was significantly higher in physically adjacent cells as compared to spatially distant cells (**Figure 4A**). From a reference list of ∼1200 ligand-receptor interactions, our analysis revealed 405 statistically significant potential communication channels (**Figure 4B, Table S5**).

**Figure 4.**
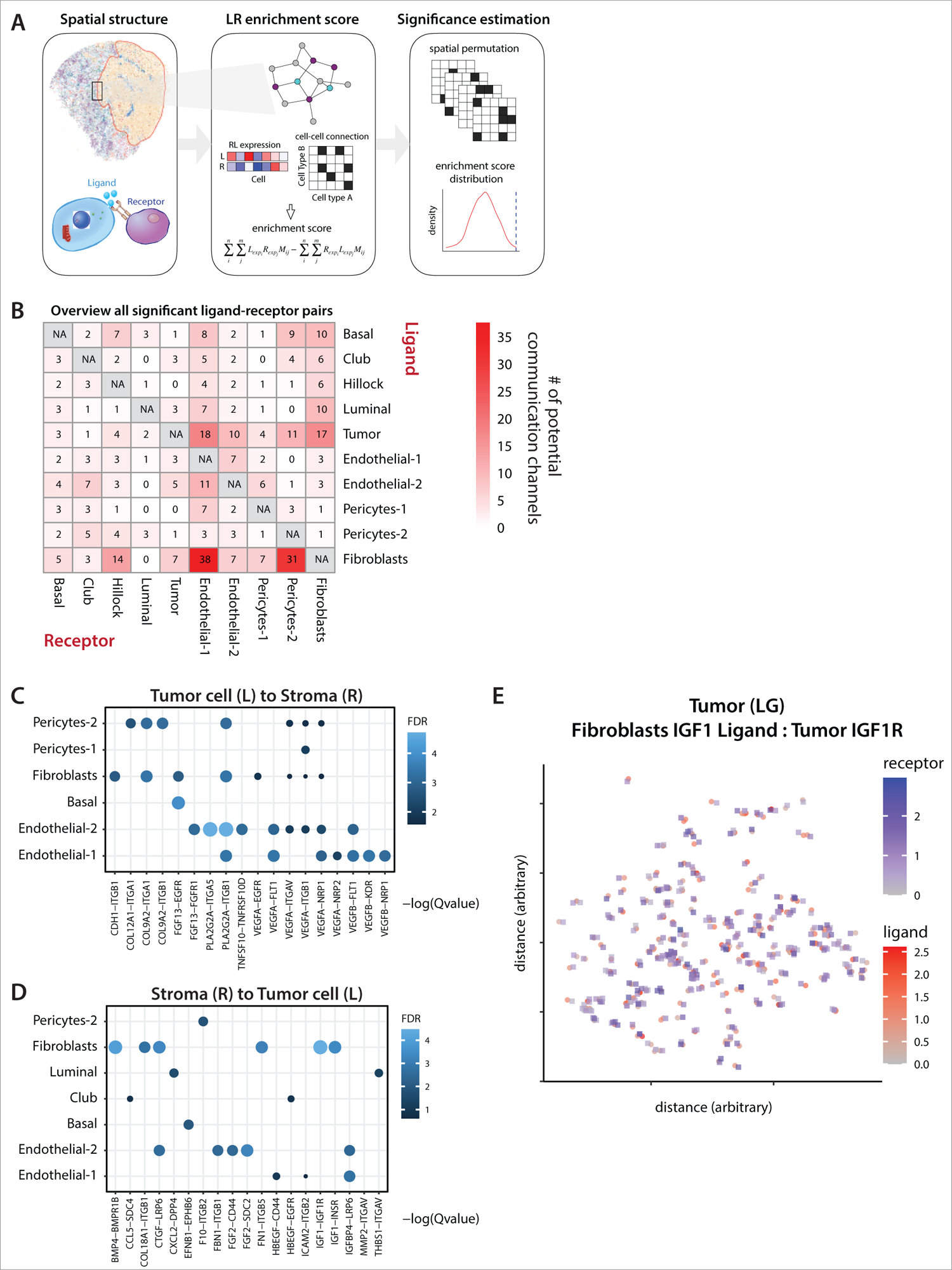
Coordination between tumor cells and stromal compartment in tumor context. **A.** Schematic of ligand receptor analysis for Slide-seq data. **B.** Summary of the total number of significant ligand–receptor interactions between stromal and epithelial cells. Each cell indicates potential channels of communication from ligand (row) to receptor (column). **C, D.** Communication channels between tumor cells and stromal cell populations, communication from tumor cells (ligand) to stromal cells (receptor) **(C**), and from stromal cells (ligand) to tumor cells (receptor) **(D**). Color and size represent significance (-log10 adjust p value) of ligand and receptor pairs, (eg Ligand IGF1 in fibroblast and receptor IGF1Rin tumor cells). **E.** Dot plot showing expression of IGF1-IGF1R axis in co-localized fibroblasts and tumor cells on a low-grade tumor case.

With a focus on tumor-stroma communication, we investigated for communication channels when considering tumor cells as a source of ligands and stromal cells as expressing receptors (**Figure 4C**). Tumor cells express vascular endothelial growth factor (VEGFA and VEGFB), which can stimulate the Endothelial-2 cells through the VEGF receptor, FLT1 (57) and beta-1 integrin (58, 59). These channels could potentially explain the pro-angiogenic shift in the state of the tumor-associated Endothelial-2 subpopulations (**Figure 3E**). We also observed potential interactions between tumor cells and fibroblasts (COL9A2-ITGA1) and tumor cells with Pericytes-2 cells (COL12A1-ITGA1), pathways that are both involved in extracellular matrix remodeling and cell migration (60–62).

Analysis of reverse interactions (i.e., stromal cells expressing ligand to a tumor receptor), revealed a potential interaction mediated by fibroblast Insulin-like Growth Factor (IGF1) stimulating tumor cell IGF1 receptor (**Figure 4D**). The IGF pathway is known to promote tumor growth and survival through suppression of apoptosis and activation of cell cycle (63). Slide-seqV2 analysis of the IGF1-IGF1R interaction confirmed the co-localization of tumor cells expressing IGF1R and fibroblasts expressing IGF1 (**Figure 4E**).

### Prostate tumors are enriched in immunosuppressive myeloid cells

Myeloid cells support tumor progression in several cancer types, and these cells are considered one of the most clinically relevant populations to target for immune therapeutic purposes (64, 65). Unsupervised clustering revealed 7 myeloid subpopulations including 3 monocyte, 3 macrophage and 1 myeloid DC (mDC) **(Figure 5A)**. Annotation was performed based on key marker genes **(Figure 5B, Figure S5A)** and validated using published monocyte and macrophage gene signatures **(Table S4, Figure 5C, panels 1 and 2)**.

**Figure 5.**
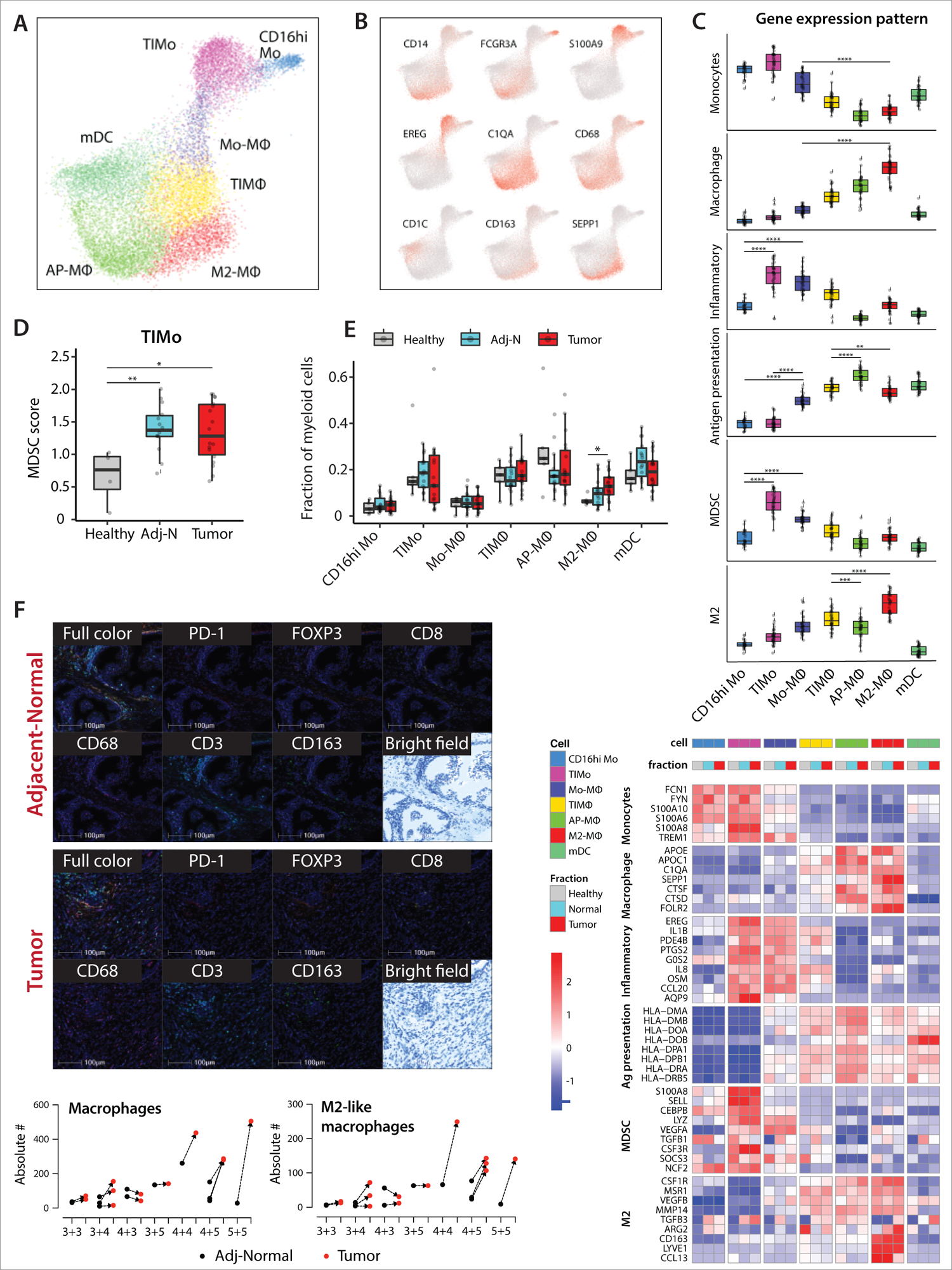
Prostate tumors are enriched in immunosuppressive myeloid cells. **A-B**. Joint embedding showing the detailed annotation of the myeloid subpopulations (**A**) and the expression of select gene markers for each myeloid subpopulation (**B**). **C**. From the top: boxplots representing the average gene expression pattern of monocyte, macrophage, inflammatory, antigen processing and presentation, MDSC gene signatures, and M2-like macrophages across the different myeloid subpopulations. Heatmap showing the average gene expression of representative genes from monocyte, macrophage, inflammatory, antigen processing and presentation, MDSC gene signatures, and M2-like macrophages gene signature across the different myeloid subpopulations in healthy, adj-normal and tumor prostate samples. See Supplementary Table S4 for the genes defining the above-mentioned signatures. **D.** Boxplot comparing the average expression of MDSC gene signature in tumor inflammatory monocytes (TIMo) across the three different samples. **E**. Boxplot representing the cell fraction of different myeloid subpopulations across the healthy, tumor and their adj-normal prostate tissues. Boxplots in **(C, D, E)** include centerline, median; box limits, upper and lower quartiles; and whiskers are highest and lowest values no greater than 1.5x interquartile range. Statistical significance was accessed using Wilcoxon rank sum test (*p<0.05, ****p<0.0001). **F.** Top: Multiplex Fluorescence immunohistochemistry (mFIHC) staining of prostate tumor tissue (bottom) and its adj-normal tissue (top) collected from a prostatectomy case of Gleason score 5+5. Samples are labeled with PD-1 (Clone EH33) (color Red), FOXP3 (color Orange), CD8 (color Yellow), CD68 (color Magenta), CD3 (color Cyan), CD163 (color Green) and DAPI (Blue) by using mFIHC. Bottom: Quantification of absolute number of macrophages (left) and M2-like macrophages (right) from mIHC data comparing tumor tissues to their matched adj-normal tissues collected from prostatectomy cases of different Gleason scores.

Monocyte subpopulations were characterized as CD16hi (CD16hi Mo) which are known as non-classical monocytes, and tumor inflammatory monocytes (TIMo) which had high expression of CD14 (**Figure 5B**, a classical monocyte marker) as well as the highest expression of an inflammatory gene signature (**Table S4, Figure 5C**). The third subpopulation was annotated as Monocyte-Macrophage (Mo-MΦ) as it showed a gradual shift in their gene expression from genes highly expressed in monocytes (e.g., S100A9) to genes expressed in macrophages (e.g., C1QA) (**Figure 5A and S5B**), suggesting a transitional cell state from monocytes toward macrophages.

Both tumor and stromal cells produce chemokines involved in the myeloid differentiation process, as well as in the recruitment of monocytes to the tumor (66). We observed high expression of CXCL12 in fibroblasts, CCL2 in pericytes and CCL3,4, and 5 in epithelial and tumor cells **(Figure S5C),** suggesting a potential role of fibroblasts and pericytes in recruiting monocytes to the prostate tumor.

Patients with prostate cancer have an ineffective immune response against the tumor and an immunosuppressive TME associated with the accumulation of myeloid derived suppressor cells (MDSCs) (67, 68). TIMo cells scored highest for an MDSC gene signature (69) **(Table S4, Figure 5C, panel 5),** and the gene signature was significantly higher in cells collected from cancerous prostate (tumor and adj-normal) compared to healthy prostate tissues **(Figure 5D)**. This suggests a role for the TIMo subpopulation in prostate tumor growth through immunosuppressive activity and the release of pro-inflammatory cytokines.

Several macrophage subpopulations were identified **(Figure 5A)**, including tumor inflammatory macrophages (TIMΦ) with a high “Inflammatory gene signature”, antigen presenting macrophages (AP MΦ) with a high “antigen processing and presentation gene signature”, as well as M2-macrophages (M2-MΦ) with a high “M2-like gene signature” **(Figure 5C, S5D, Table S4)**. M2-MΦ showed a gradual increase in cell abundance from healthy towards tumor fraction **(Figure 5E)** and M2-like macrophages have been shown to suppress anti-tumor immune response across a broad range of tumors (70). In prostate cancer, the high infiltration of M2-like macrophages in tumor tissue has been linked to tumor recurrence (71) and metastasis (72, 73).

Multiplex immunohistochemistry (mIHC), performed *in-situ* on the same tissue samples as the single cell expression, confirmed a higher infiltration of CD68+ macrophages and of CD68+CD163+ M2-MΦ in tumor tissues compared to their matched adj-normal tissues **(Figure 5F)**. Quantification of tumor infiltration by M2-MΦ was more pronounced in cases of high Gleason scores (4+4, 4+5, 5+5) (**Figure 5G)**. M2-MΦ express high levels of genes involved in angiogenesis such as angiogenic factor EGFL7 (74, 75) and in tumor metastasis such as LYVE1 (76, 77) and NRP1 (78) **(Figure S5E)**, suggesting a role for M2-MΦ infiltration in angiogenesis within tumors.

Myeloid dendritic cells (mDCs) present tumor antigens to T-cells with a critical role in the initiation and regulation of the adaptive anti-tumor immune response (79–81). We identified three mDC subpopulations, each with high expression of either CD1C, CLEC9A or LAMP3. No significant changes were observed in the cell abundance of the different mDCs subsets (**Figure S5F**). Overall, our myeloid cell analysis identified immunosuppressive subpopulations that may contribute to tumor progression, including MDSC-like monocytes (TIMo), and macrophages with an M2-like signature.

### Prostate cancer is characterized by T-cell exhaustion and immunosuppressive Treg activity

The adaptive immune system plays a pivotal role in mounting an effective, antigen-specific immune response against tumors. Unsupervised clustering of the lymphoid compartment revealed four CD4+ T cell, three CD8+ T cell and two NK subpopulations **(Figure 6A)** as annotated by key-marker genes (**Figure 6B)**.

**Figure 6.**
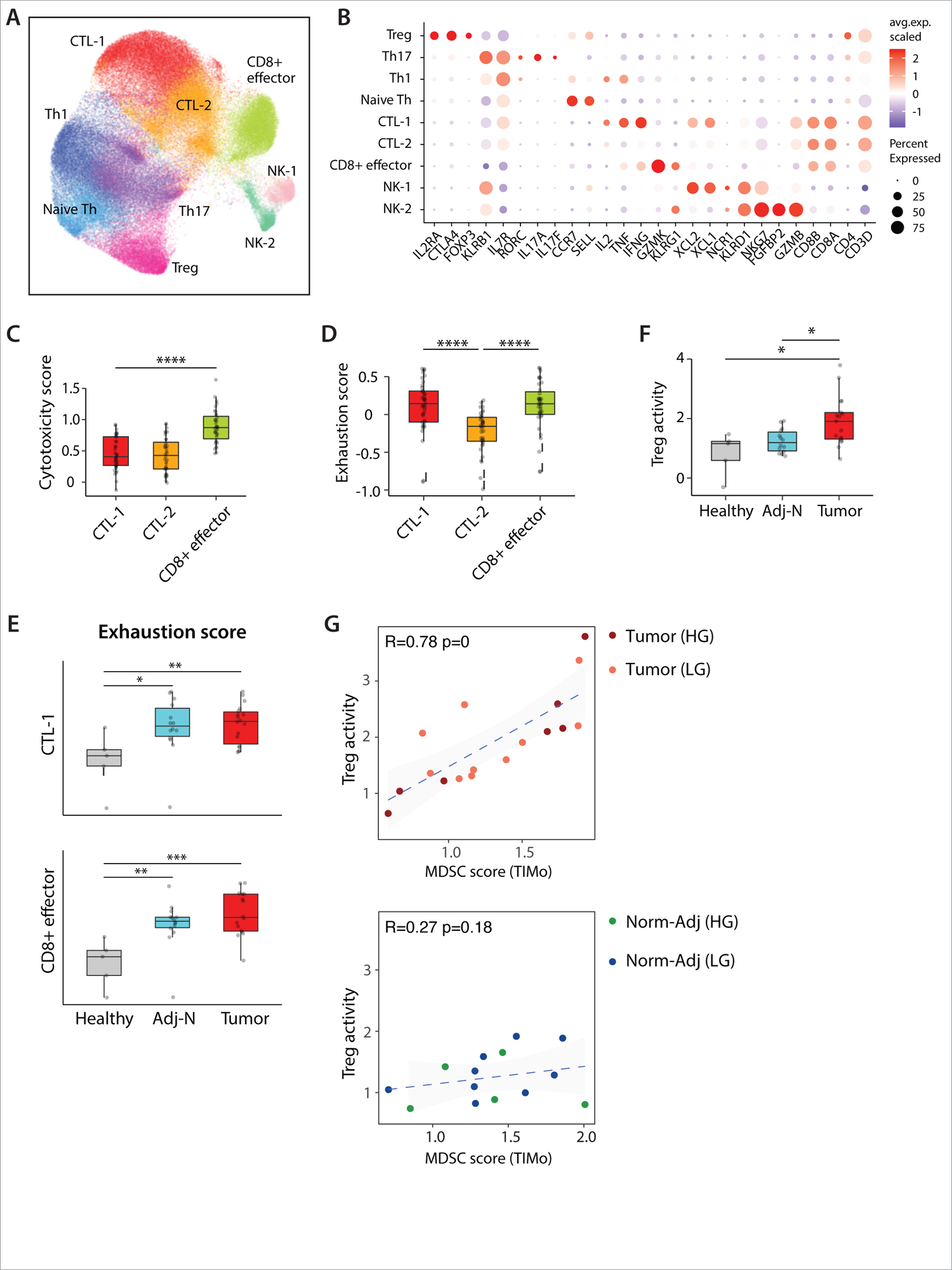
Prostate cancer is characterized by T-cell exhaustion and immunosuppressive Treg activity. **A.** Joint embedding showing the detailed annotation of lymphoid subpopulations. **B**. Dotplot representing key-marker gene expression in lymphoid subpopulations. The color represents scaled average expression of marker genes in each subpopulation, and the size indicates the proportion of cells expressing marker genes. **C-D**. Boxplots represent the average expression of cytotoxicity **(C)** and exhaustion **(D)** scores in CD8+ CTLs subpopulations (CTL-1, CTL-2 and CD8+ effector cells). **E**. Boxplots comparing the average expression of exhaustion score in CTL-1 (top) and CD8+ effector (bottom) subpopulations across healthy, adj-normal and tumor samples. **F**. Boxplot represents the average expression of Treg activity gene signature in Treg subpopulation across the three different samples. Boxplots in **(C-F)** include centerline, median; box limits, upper and lower quartiles; and whiskers are highest and lowest values no greater than 1.5x interquartile range. Statistics significance are accessed using Wilcoxon rank sum test.(*p<0.05, **p<0.01, ****p<0.0001). **G**. Scatter plot showing the correlation between Treg activity in Tregs and MDSC score in TIMo subpopulation in tumor (top) and adj-normal prostate tissues (bottom). Each dot represents a sample.

The functional state of CD8+ T cells was assayed using a cytotoxicity gene signature (“cytotoxicity score”) **(Table S4)** (82, 83). CD8+ effector cells exhibited a higher cytotoxicity score compared to the other CTL-1 and CTL-2 CD8+ subpopulations **(Figure 6C)** and the CD8+ effector cell cytotoxicity score was consistent across different sample fractions **(Figure S6A)**.

Both CTL-1 and CD8+ effector cells exhibited higher expression of a T-cell exhaustion gene signature (3,84,85) **(Table S4, Figure 6D)**, and the exhaustion score was higher in the prostate tumor and adj-normal samples as compared to healthy prostate tissues **(Figure 6E)**. No significant difference in the exhaustion score was observed when comparing cells from LG and HG samples **(Figure S6B)**.

Measurement of T cell abundance showed a higher proportion of exhausted CTL-1 cells in tissues collected from cancerous prostate compared to healthy prostate tissues **(Figure S6C),** suggesting an expansion of exhausted CTLs in the prostate tumor. No differences were observed in T cell abundance when comparing LG and HG Gleason groups **(Figure S6D).**

CD4+ T cells were subdivided into naïve, T-helper-1 (Th1), T-helper 17 (Th17), and T-regulatory (Treg) cells based on cell-type specific genes (86) **(Figure 6B)**. CD4+ cell abundance was stable across the different sample fractions **(Figure S6C and S6D).** As a surrogate for CD4+ T-cell function, Treg activity was assayed (87, 88) (**Table S4**) and was increased in the tumor and adj-normal samples **(Figure 6F)**. Notably, genes of tumor necrosis factor receptor superfamily TNFRSF9, TNFRSF18, and TNFRSF4 were highly and exclusively expressed in the Tregs infiltrating the tumor **(Figure S6E)**. These receptors bind tumor necrosis factors, pro-inflammatory cytokines involved in inflammation-associated carcinogenesis (89) and in supporting an immunosuppressive TME.

Tregs and MDSCs represent two immunosuppressive cell populations important for cancer immune tolerance. Both populations exhibited high suppressive activity in the tumor fraction and their crosstalk has been previously reported in different cancers (90, 91). Based on this, we examined the correlation between the MDSC score in TIMo and the Treg activity score in Tregs both in tumor samples and their adjacent-normal tissue samples. Within the tumor fraction, the MDSC score and Treg activity score were significant correlated, with no clear separation between LG and HG Gleason patients **(Fig 6G, top).** No correlation was seen in adj-normal tissues **(Fig 6G, bottom)**.

Taken together, we characterized the functional status of T-cell subpopulations in prostate tumors to demonstrate exhausted CTLs along with increased Treg suppressive activity which correlated strongly with the suppressive activity of MDSC-like monocytes.

### The prostate cancer TME is enriched in exhausted CD56<u>^DIM^</u> NK cells

Natural killer (NK) cells are an innate lymphoid cell with cytotoxic function that can be modulated by activating and inhibitory cell-surface receptors (92). A high density of tumor infiltrating NK cells usually correlates with good prognosis in different solid tumors, including breast cancer (93), lung cancer (94), and prostate cancer (95). NK cells were annotated based on key marker gene expression **(Figure 6B)** (96) and clustering revealed 4 NK subpopulations **(Figure 7A and 7B).** No differences were observed in NK cell abundance across the 5 different sample fractions **(Figure S7A).** NKT cells were characterized by high expression of T cell marker genes CD3D and CD8 and CD56dim NK cells by high expression of HAVCR2, which is expressed by terminally differentiated NK cells (96). CD56bright NK cells expressed XCL1, XCL2, GZMK, CD44 and KLRC1 (96), while the CD56bright-IL7R+ cells separated based on specific expression of IL7R and the homing-receptor SELL (encoding CD62L) **(Figure 7B)** (97–99).

**Figure 7.**
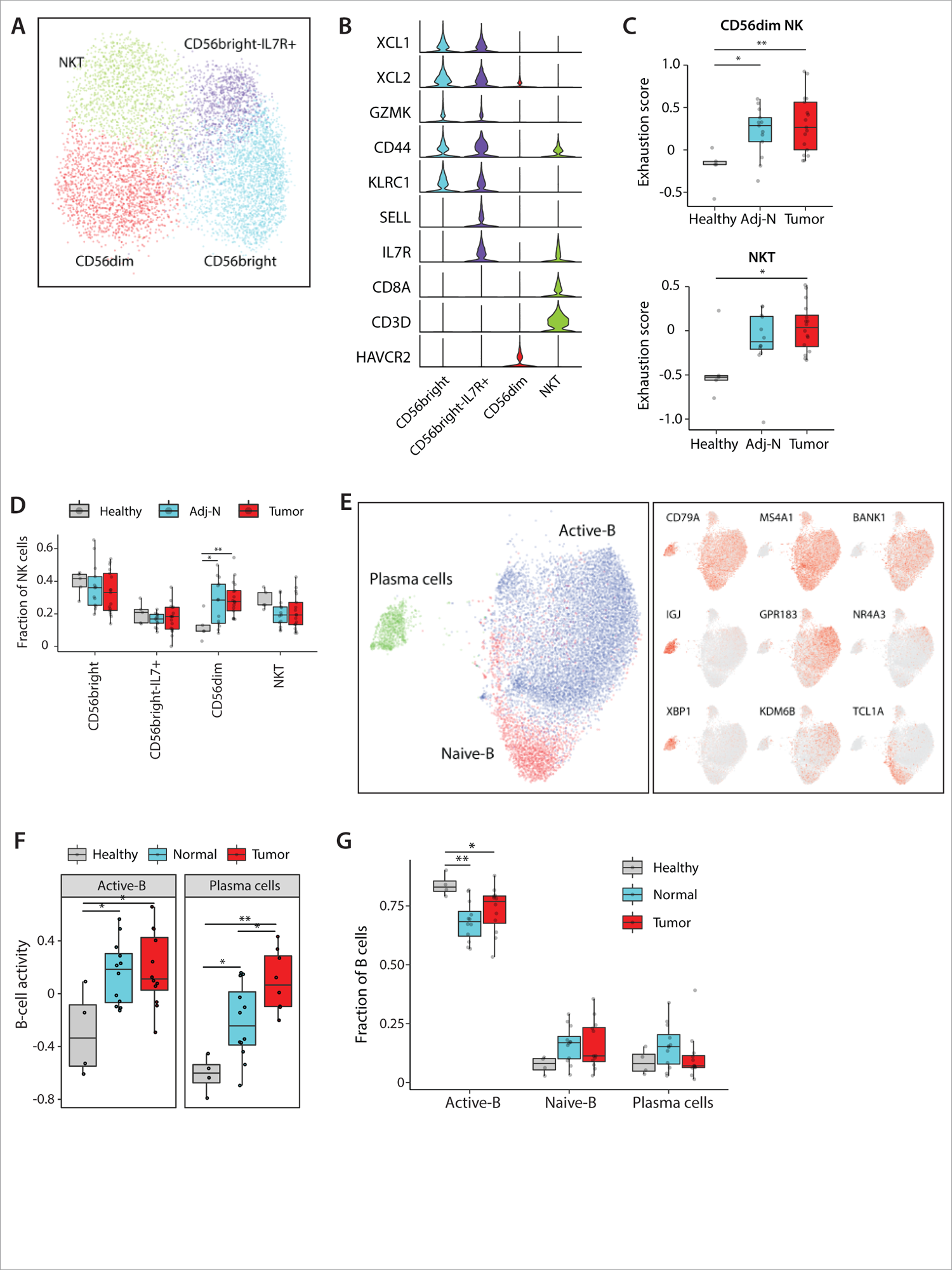
The prostate cancer TME is enriched ins characterized by exhausted CD56D^IM^ NK cells and activated B cells. **A.** Joint embedding showing the detailed annotation of NK subpopulations. **B.** Violin plot showing the average expression of indicated marker genes in NK subpopulations. **C**. Boxplots comparing the exhaustion score of CD56dim and NKT subpopulation across healthy, adj-normal, and tumor samples. See Supplementary Table S4 for the genes defining exhaustion score. **D**. Boxplot comparing the relative abundance of different NK subpopulations in healthy, adj-normal, and tumor samples. **E**. Joint embedding showing the detailed annotation of B cell subpopulations (left) and the expression of B cell specific marker genes (right). **F**. Boxplot comparing B cell activity signature in active B and plasma subpopulations between healthy, adj-normal, and tumor samples. **G**. Boxplot comparing the relative abundance of each B subpopulations across healthy, adj-normal and tumor samples. Boxplots in **(C-D,F-G)** include centerline, median; box limits, upper and lower quartiles; and whiskers are highest and lowest values no greater than 1.5x interquartile range. Statistical significance was accessed using Wilcoxon rank sum test (*p<0.05, **p<0.01).

The NKT and CD56^DIM^ cells also showed high expression of the effector protein and cytotoxic-related genes FGFBP2, GNLY, GZMB, GZMH (96, 100) **(Figure S7B)**. However, these same NK subpopulations exhibited a higher exhaustion gene signature (**Table S4**) in the tumor samples as compared to healthy tissue (**Figure 7C**), suggesting impaired effector function within the prostate TME. Of the NK subpopulations, the CD56^DIM^ cells scored highest for the exhaustion gene signature (**Figure S7C**) and were in higher abundance in the prostate tumor as compared to healthy prostate (Figure 7D).

### The prostate cancer TME is characterized by activated B cells

B cells are less extensively studied in cancer as compared to the myeloid and T cell counterparts. B cell infiltration has been described in several cancer types though their function and correlation to survival remain controversial (101). Clustering of B cells based on key marker genes revealed 3 subpopulations: naïve-B, active-B and plasma cells **(Figure 7E)**. B-cell abundance was similar across the five sample fractions (**Figure S7D**). B cell activity was assessed in active B cells and plasma cells (102) (**Table S4**). B-cell activity was significantly higher in cells from tumor and adj-normal tissue compared to healthy prostate **(Figure 7F)**, possibly due to the recognition of tumor antigens by the B-cells. However, this increased activity was accompanied by a lower B-cell abundance in the tumor samples **(Figure 7G)**.

In our spatial characterization of immune cells, B cells and macrophages were most abundant, with few monocytes, T cells and plasma cells (**Figure S7E, S7F and S7G**). This low abundance did not permit a formal analysis of potential ligand/receptor interactions.

## Discussion

Localized prostate cancer has been extensively studied using bulk transcriptomic and genomic sequencing approaches, providing insights into oncogenic drivers and recurrent molecular changes. Here, we used a high-resolution single-cell approach to characterize changes in tumor, immune, and non-immune stromal cells within the with tumor microenvironment. These findings were complemented by spatial transcriptomic analysis where the tissue architecture and cell-to-cell relationships are preserved, allowing one to determine whether transcriptomic changes are context-dependent.

The strengths of our study include the (a) fresh nature of our patient samples, (b) matched tumor and adjacent-normal samples across a spectrum of Gleason scores to help overcome the inherent patient-to-patient variability, (c) rigorous collection of truly normal control prostate samples (healthy), and (d) the combined single-cell and spatial transcriptomic analysis. Indeed, this manuscript represents a highly detailed spatial transcriptomic analysis using Slide-seqV2 to characterize the prostate tumor tissue, as well as a new computational approach to detect spatial context-dependent transcriptional differences in different cell types, which are typically obscured by the admixture from neighboring cells. Such changes are likely to provide insights about the impact of microenvironment on the cell and the mechanisms through which such changes may be induced. We hope that the developed context-dependent DE method and the associated tutorial will enable analysis of such processes by other investigators (**Supplementary Note 1**).

As expected, the prostate TME is complex with several subsets of myeloid cells, T cells, NK cells and B cells in addition to the non-immune stromal populations of endothelial cells, fibroblasts and pericytes. This led to some key observations.

Regarding epithelial cells, we identified distinct subsets of hillock and club cells that have been described in normal prostate tissue (5,7,20). Club cells have been identified in human prostate tumors (23); however, we are the first to show and characterize the hillock cells in human prostate tumors (**Figure 2B**). Our RNA velocity analysis suggested a progenitor role for club cells which has been previously reported (23). We also saw a directional flow from hillock to basal cells, suggesting a second progenitor role of the hillock cells. The identification of hillock epithelial subset in our dataset may be due to the dissociation protocol we followed as it has been reported that the method and conditions of tumor dissociation affects cell yield and transcriptional state in primary solid tumor tissues (103, 104). However, hillock epithelial cells were also detected in our Slide-seq data where no dissociation took place (**Figure 2E**).

Malignant and normal cells can be challenging to distinguish. We used an iterative strategy, first relying on detection of genomic aberrations to distinguish normal and malignant luminal-type cells, and then deriving a succinct Prostate Tumor Gene Signature, which could robustly identify tumor cells across four independent datasets.

Regarding myeloid cells, we showed that a population of tumor-inflammatory monocytes were immunosuppressive with a high MDSC gene signature. In addition, M2-like macrophages were increased in abundance in the tumor microenvironment, a finding that was consistent across single-cell analysis and immunohistochemistry. M2-macrophages have been reported to be involved in the growth and progression of prostate cancer and they have gained remarkable importance as therapeutic candidates for solid tumors (105).

Regarding lymphoid cells, we observed that cytotoxic T-lymphocytes showed a high exhaustion signature along with a low cytotoxic signature. Treg cells also showed a high exhaustion signature. Interestingly, we did not see significant T-cell differences when comparing low-grade and high-grade cases, suggesting that even the low-grade tumors had already established a highly immunosuppressive microenvironment. Even within the NK cells, the CD56^DIM^ NK cells were expanded in the tumor fraction, again suggesting a functionally less cytotoxic NK cell.

We hypothesized that the immunosuppressive myeloid cells were contributing to the exhausted T-cell phenotype, as our group has previously shown in the setting of metastatic prostate cancer (106). Indeed, there was a correlation between the MDSC and Treg activity signatures, pointing to the role of myeloid cells in establishing a T-cell suppressive and pro-tumor microenvironment.

We utilized the spatial neighborhood to infer cell-to-cell interactions with high resolution and this enabled the identification of ligand-receptor interactions in undissociated tissue section, especially between tumors cells and their stroma. Beyond the tumor-fibroblast and tumor-endothelial cell communication that we highlighted we hope that this analysis will prove more broadly useful for the community and point towards clinically relevant and therapeutically targetable interactions. This analysis also supports the complementary use of techniques that involve tissue dissociation with techniques that preserve the normal tissue architecture to home in on these cell-cell relationships.

Overall, this combined dataset of single-cell and spatial transcriptomic analysis of primary prostate tumor samples and their normal controls provides a rich community resource. Biological validation of the tumor relationships with their neighboring immune and stromal cells will lead to a better understanding of prostate cancer progression and will identify new therapeutic targets for this common disease. We also hope that this manuscript highlights the importance of multidisciplinary teams as the longitudinal collection of fresh patient samples can only be obtained when surgical, pathology, and basic science collages work in true collaboration.

## STAR Methods

### Patient materials

In accordance with the U.S. Common Rule and after Institutional Review Board (IRB) approval, all human tissues ware collected at Massachusetts General Hospital in Boston (MGH, Boston, MA) and carried out with institutional review board (IRB) approval (IRB#2003P000641).

### Surgical approach and tumor collection

Patients with clinically localized prostate cancer were treated with minimally invasive transabdominal radical prostatectomy. The dissection of the prostate was done by antegrade approach, freeing the bladder neck, then progressing caudally to the apex and urethra. Upon freeing the prostate, it was placed in a laparoscopic specimen sac. The specimen was then immediately removed from the patient. The staff transported the tissue without delay to the pathology lab where the research staff was waiting to assure the least possible ischemic time from separation of the organ from blood supply to prepared specimen. The prostate was marked with ink, and sectioned. The prostate cancer tissue is identified by a trained genitourinary pathologist, aided with biopsy and MRI reports. The cancer is confirmed by histological examination of the immediate adjacent tissue. Cancer cell content is estimated to be 70%.

### Sample preparation

*Dissociation of tissues into single cells:* All samples were collected in Media 199 supplemented with 2% (v/v) FBS. Single cell suspensions of the tumors were obtained by cutting the tumor in to small pieces (1mm^3^) followed by enzymatic dissociation for 45 minutes at 37°C with shaking at 120 rpm using Collagenase I, Collagenase II, Collagenase III, Collagenase IV (all at a concentration of 1mg/ml) and Dispase (2mg/ml) in the presence of RNase inhibitors (RNasin (Promega), RNase OUT (Invitrogen)), and DNase I (ThermoFisher). Erythrocytes were subsequently removed by ACK Lysing buffer (Quality Biological) and cells resuspended in Media 199 supplemented with 2% (v/v) FBS for further analysis.

*FACS sorting:* Single cells from tumor samples were surface stained with anti-CD235-PE (Biolegend) for 30 min at 4°C. Cells were washed twice with 2% FBS-PBS (v/v) followed by DAPI staining (1 ug/ml). Flow sorting for live-nonerythroid cells (DAPI-neg/CD235-neg) was performed on a BD FACS Aria III instrument equipped with a 100um nozzle (BD Biosciences, San Jose, CA). All flow cytometry data were analyzed using FlowJo software (Treestar, San Carlos, CA).

### Multiplex immunohistochemistry analysis

We used multiplex immunohistochemistry (mIHC) panel to evaluate a set of unselected radical prostatectomy cases, spanning all grade groups. A seven-plex Fluorescence Immunohistochemistry assay was performed on 4-µm FFPE sections, using Leica Bond Rx autostainer. A six antibodies panel consisted of CD3 (Rabbit polyclonal, Dako), CD8 (C8/144B, Mouse monoclonal, Dako), PD-1(EH33, Mouse monoclonal, Cell Signaling), FOXP3 (D2W8E, Rabbit monoclonal, Cell Signaling), CD68 (PG-M1, Mouse monoclonal, Dako), CD163 (10D6, Mouse monoclonal, Leica Biosystem), along with DAPI counterstaining. Briefly the staining consists of sequential tyramine signal amplified immunofluorescence labels for each target, and a DAPI counterstain. Each labeling cycle consists of application of a primary antibody, a secondary antibody conjugated to horse radish peroxidase (HRP), and an opal fluorophore (Opal 690, Opal 570, Opal 540, Opal 620, Opal 650 and Opal 520, Akoya Biosciences), respectively. The stained slides were scanned on a Perkin Elmer Vectra 3 imaging system (Akoya Biosciences) and analyzed using Halo Image Analysis platform (Indica Labs). Each single stained control slide is imaged with the established exposure time for creating the spectral library. We ran an algorithm learning tool utilizing the Halo image software training for the gland and stroma regions, and subsequently completed cell segmentation. The thresholds for the antibodies were set respectively, based on the staining intensity, by cross reviewing more than 20 images. Cells with the intensity above the setting threshold were defined as positive. Regions of interest included both immune-cell-rich and non-rich areas and included both tumor and benign areas.

### scRNA-seq data processing and analysis

Sequencing data were processed using 10X Cell Ranger with default parameters (version 3.0.1), aligned to GRCh37 human reference genome. The obtained read count matrices were further analyzed with scrublet (107) for doublets identification. Scrublet scores above 0.4 were omitted from further analysis. In total, 179,359 cells from 39 samples were obtained. We used Conos (15) (k=15, k.self=5, matching.method=’mNN’, metric=’angular’, space=’PCA’) to integrate multiple scRNA-seq datasets together. Principal component analysis was performed on 2000 genes with the most variable expression was selected by conos. Leiden clustering was used to build to determine joint cell clusters across the entire dataset collection. First 15 principal components were used to perform UMAP embedding.

### Determination of major cell types and cell states

To identity major cell types in both tumor sample datasets and healthy sample datasets, we used sets of well-established marker genes for each of those cell types and annotated each cell type based on highly expressed genes. The detailed gene list can be found in **Table S3**. For subtype assessment within the major cell types, we extract raw count matrices and re-analyzed cell subsets separately with Conos.

### Calculation of gene set signature scores

To assess cell states in different cell subsets and conditions, we used a gene set signature score to measure the relative difference of cell states. The signature scores were calculated as average expression values of the genes in a given set. Specifically, we first calculated signature score for each cell as an average normalized (for cell size) gene expression magnitudes, then the signature score for each sample was computed as the mean across all cells. All signature gene modules are listed in the Supplementary Table S4. The statistical significance was assessed using Wilcoxon rank-sum test.

### Differential expressed genes (DEG) analysis

For differential expression analysis between cell types, Wilcoxon rank sum test, implemented by the getDifferentialGenes() function from Conos R was used to identify marker genes of each cell cluster. The genes were considered differentially expressed if the p-value determined Z score was greater than 3. For differential expression analysis between sample fractions (for example Tumor Treg vs. adj-Normal Treg), getPerCellTypeDE() function in Conos was utilized with default settings. DESeq2 (108)was applied to “mini-bulk” (or meta-cell) RNA-seq measurements by combining all molecules measured for each gene in each subpopulation in each sample. A minimal number of 10 cells (of the selected cell type) were required for a sample to be included in the comparison.

### Identification of tumor cells from luminal epithelial cells

To identify the tumor cells from normal epithelial cells, we used inferCNV cell (24, 25) to inferred large-scale chromosomal copy-number variations. We performed inferCNV on different epithelial subpopulations using the same cell type from healthy tissues as the reference “normal” cells. Only epithelial liminal cells show clear CNV aberration. To identify tumor cells, we examinated cell hierarchical clustering of CNVs profile obtained from inferCNV and filter tumor cells with deletion in chr8, chr12 and chr16. In addition, “prostate cancer signature” below were used to rescue additional tumor cells. In total, 1,237 tumor cells were obtained.

### Generation of the “Prostate Tumor Gene Signature”

To generate a gene expression signature that is clinically applicable, we compared the gene expression profiles between tumor cells and non-tumor luminal cells in tumor fraction. Only the upregulated genes with an Zscore > 3 were selected and taken into subsequent analysis. We next screened each of the DEGs based on their expression in healthy prostate tissue, requiring gene expressed less than 5% cells of all epithelial cells. In total, we identified 8 significant DEGs met the above criteria. After that, we defined the eight-gene set diagnosis signature score, calculated as average expression values of the eight gene set. We applied the gene diagnosis signature score to multiple bulk RNAseq data to quantify the predictive accuracy of the diagnosis-related DEGs. ROC analysis showed a strong prostate cancer predictive ability with an AUC score of 0.956 (GSE21034 (27)), 0.93 (GSE97284 (28)), 0.937 (TCGA (29)) and 0.94 (GSE70770 (30)) in four independent prostate cancer cohorts.

### RNA velocity-based cell fate tracing

To perform the RNA velocity analysis, the spliced reads and unspliced reads were recounted by the velocyto python package (21) based on previous aligned bam files of scRNA-seq data. The calculation of RNA velocity on low diminutions UMPA embedding were done by following the scvelo python pipeline scVelo (22) was applied to both merged datasets and individual sample group.

### Slide-seq data pre-processing and cell-type annotation

Sequencing data were processed using Slide-seq tools pipeline (https://github.com/MacoskoLab/slideseq-tools), aligned to human genome reference version hg38. After obtained count matrixes and beads spatial coordinates, RCTD (17) was used to annotated barcoded beads. Specifically, we sample down 10X scRNA-seq data to 1,000 cells per cell type and import 10X data into the RCTD object as reference. Slide-seq data were filtered using default RCTD setting, requiring at least 100 UMI per cell. To annotate Slide-seq beads, We first annotated the major cell clusters (T cells, B cells, stromal cells, epithelial cells and myeloid cells) with corresponding 10X reference in major cell annotation (run.RCTD), then each of major cell cluster was extracted for cell sub-cluster annotation. We only keep result in singlet and doublet certain prediction categories.

### Spatial autocorrelation analysis

To measure how cells are spatially distributed in context, we used spatial autocorrelation to evaluate clustering centrality pattern of each cell type. We applied autocorrelations function from hotspot package (109), and furthermore Moran’s I was used to measures the overall spatial autocorrelation of Slide-seqV2 data. Positive value indicates the centralized clustering. We then used willcoxon test to access Moran’s I differences between healthy, adjacent normal and tumor fractions.

### Estimate spatially differential expressed genes

To obtain the differentially expressed genes across different regions within a puck we used a custom pre-processing phase. We first identified specific regions within the tumor puck by segmenting out the tumor proliferated region as “tumor context” and the non-proliferated region within the puck as “tumor-adjacent context”. The context specific cell level expressions are then summarized to the cell-type level pseudo-bulk profiles. We use a constrained linear regression model to correct for the ad-mixture effects in the slide-seqV2 measurement given a target cell-type. Finally, we pass the corrected pseudo-bulk profiles to the off-the-shelf differential gene expression tool edgeR (110). For the detailed overview of the differential expression pipeline please refer to the Supplementary Note.

### Identification of significant ligand-receptor pairs

The ligand-receptor (LR) pairs were downloaded from published databases CellPhoneDB (v1.1.0) (111). To identify significant ligand-receptor pairs in 10X data, we use a similar approach as previously described (111). We first calculate gene expression ratio in each cell type and only considered genes with more than 10% of cells demonstrating expression within each cell type. We then calculated average expression of ligand and receptor pairs across cell type pairs in normalized scRNA-seq data. The product of average ligand expression in cell type A and the average receptor expression in cell type B was used to measure LR pair expression. Statistical significance was then assessed by randomly shuffling the cluster labels of all cell types and re-calculating ligand-receptor average pair expression across 1,000 permutations, which generated a null distribution for each LR pair in each pairwise comparison between two cell types. P-values were calculated with the normal distribution curve generated from the permuted LR pair interaction scores.

To access ligand-receptor interactions in slide-seq data, we combined information from cell spatial structure and ligand-receptor expression. We assume that spatially inferred ligand-receptor pairs are co-expressed in adjacent cells. Specifically, we first build a nearest neighbor graph (mNN) based on beads spatial coordinates, then for any pair of cell types, we defined a ligand-receptor (LR) score to filter informed ligand-receptor pairs by calculating the aggregated expression product of ligand and receptor in adjacent neighborhood cells obtained from mNN graph. LR score is defined as:

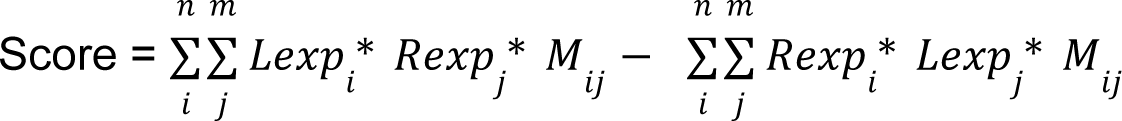

Where n represents the number of cells for “sender cell” type A, m represents the number of “receiver cells” for cell type B. *Lexp_i_* represents Ligand L expression in cell type *A_i_*. *Rexp_j_* represents Receptor R expression in cell type *B_j_*. *M_ij_* is cell adjacency matrix for cell type A and B. To avoid potential bias from admixture noise, like ligand expression signal from “receiver cells” B and receptor expression signal from “sender cell” A, We defined a revers score with switched ligand and receptor. *Rexp_i_* represents Receptor R expression in cell type *A_i_*. *Lexp_j_* represents Ligand L expression in cell type *B_j_*. To evaluate if the RL score S is statistically significant, we created a background distribution by shuffling cell labels in expression matrix for 2000 times. In each round, a permutation score is calculated using the same formula. P-values were calculated as the probability of observed RL score in background distribution. The p values for all ligand-receptor pairs in cell-type pairs were subsequently adjusted for multiple hypothesis testing. In total, 405 significant potential interaction were obtained shown in Table S5.

## Supporting information

Table S3-Gene-list-cell-annotation

Table S4-Gene list_ gene signatures

Table S1-Patient-clinical-characterisation

Table S5-LR-Slideseq-data

Table S2-Data quality

## Acknowledgements

We are particularly indebted to our patients and their clinical care teams. We gratefully acknowledge support from Bill & Cheryl Swanson. We acknowledge funding from NIH CA193481 and DK103074 (to DTS and PVK), National Cancer Institute CA 163191 (to DTS), NIH R01HL131768 (to PVK), European Research Council Synergy (‘KILL OR DIFFERENTIATE’, 856529, ERC-2019-SyG) to PVK, Dana-Farber / Harvard Cancer Center Nodal Award (CCSG grant P30CA006516), the Harvard Ludwig Cancer Center, the Harvard Stem Cell Institute and the Gerald and Darlene Jordan Professor of Medicine Chair (to DTS). N.B. was funded by the Swedish Cancer Society and Swedish Childhood Cancer Fund. Y.K was supported by a grant from the STARR cancer consortium. K.S. was funded by the Urology Care Foundation Research Scholar Award and Prostate Cancer Foundation Young Investigator Award. Olga Kharchenko designed the medical illustration in Figure 1A and the graphical abstract. Patient samples were sorted at the HSCI/CRM flow cytometry core facility at MGH.

## Author contributions

T.H., N.B., D.M.D., D.B.S., and P.J.S. conceived the study. P.J.S. coordinated the multi-disciplinary teams and the IRB-approved protocol. T.H., S.M., N.B., P.J.S., D.B.S., and P.V.K. directed the study. Sample collection methodology and surgeries were performed by D.M.D. D.Z. and M.W. provided the healthy prostate tissues from cystoprostatectomy cases. D.M.D. provided the prostate tissues from prostatectomy cases. Human samples were collected and isolated, and libraries prepared by T.H., N.B., and Y.K. Slide-seq arrays and library preparation were performed by E.M. in the labs of F.C. and E.Z.M. Slide-seq was obtained at F.C and E.Z.M labs at the broad. S.M., H.S., and P.V.K. performed the computational analysis. T.H., S.M., H.S., N.B., D.B.S. and P.V.K. interpreted the data. T.H., S.M., H.S., D.B.S. and P.V.K. wrote the manuscript. All authors read, edited, and approved the manuscript.

## Data and code availability

The accession numbers for the raw sequencing data and processed data in this paper are under the accession number: GSEXXXXXX^1^. Custom code that was used in this study can be found on github at https://github.com/shenglinmei/ProstateCancerAnalysis. The differential expression tool for Slide-seqV2 is available at https://github.com/kharchenkolab/slideseqde.

**Figure S1. The prostate TME characterized by single-cell and spatial transcriptomic analysis. A.** Representative photomicrographs of matched adjacent-normal prostatic tissue (A1 and B1, 4x) and prostate cancer (PCa) (A2: Gleason score 4+3 and B2: Gleason score 3+5, 20x) from radical prostatectomy specimens of two PCa patients and healthy prostate tissue from cystoprostatectomy specimen of a bladder cancer patient (C1, 4x). **B**. Heatmap shows an overview of marker genes (row) expressed in major cell populations (column). **C.** UMAP embedding showing major cell populations obtained using two different dissociation protocols on an adj-normal prostate tissue (Gl 3+3) separately (left: Collagenase+Dispase, right: Rocky; details in Methods section). **D.** The Heatmap demonstrating spearman correlation coefficients of gene average expression level between Rocky (x-axis) and Collagenase+Dispase (y-axis) in each cell type. **E**. Barplot representing the fraction of major cell populations within each sample fraction collected for 10x. **F**. Barplot representing the fraction of major cell populations within each sample collected from 10x Genomics data. **G.** Dotplot representing key-marker gene expression in major cell types in Slide-seqV2. The color represents scaled average expression of marker genes in each cell type, and the size indicates the proportion of cells expressing marker genes.

**Figure S2. A Prostate Tumor Gene Signature distinguishes normal and malignant luminal epithelial cells. A**. Dotplot showing the average expression of select marker genes in epithelial subpopulation across healthy, adj-normal and tumor samples. The color represents scaled average expression of select marker genes in each epithelial subpopulation, and the size indicates the proportion of cells expressing select marker genes. **B.** Gene Ontology terms enriched in the top 200 high-loading DE genes of the different epithelial subpopulation comparing tumor to healthy samples. **C.** Inferred CNV profile of malignant cells and normal epithelial luminal cells from tumor sample, using epithelial luminal cells from healthy samples as the reference. **D.** ROC curves for “Prostate Tumor Gene Signature” score applied on four independent prostate cancer datasets (TCGA, GSE21034, GSE97284, GSE70770) (see method).

**Figure S3. Heterogeneity of malignant cells. A.** Heatmap showing DE genes in the three malignant cell subclusters. **B.** Overview of enriched GO terms of top 200 upregulated genes for each malignant cell subcluster compared to all malignant cells. **C.** Heatmap showing the average gene expression of EMT gene signature in malignant cells and epithelial luminal cells in healthy, adj-normal and tumor prostate samples. **D.** Comparison of spatial autocorrelation (Moran’s I) of epithelial club and epithelial Hillock in healthy, adj-normal and tumor samples. Statistical analysis was accessed using Wilcoxon rank sum test (*p<0.05, error bars: SEM). **E.** Zoomed in view of region within the tumor (HG) puck, shows the high constellation of tumor cells in the tumor-enriched region contrasting heterogeneous cell-type population on the other side.

**Figure S4. A high endothelial angiogenic activity within prostate tumor microenvironment. A**. Dotplot representing key-marker gene expression in stroma subpopulation. The color represents scaled average expression of marker genes in each cell type, and the size indicates the proportion of cells expressing marker genes. **B.** The differential expression tests are performed on two tumor replicates (two pucks corresponding to Tumor01 and Tumor02 from the same sample of Tumor (HG)). We model the changes in gene expression comparing the tumor context and the tumor-adjacent context. The scatterplots are showing the gene expression of Endothelial-2 cells (log-transformed pseudo-bulk) in tumor-adjacent context (x-axis) vs. the tumor-context (y-axis). The left scatter plot is with the gene expression values before correction and shows the tumor marker genes (in red) to be significantly differentially expressed while the right scatter plot is with the corrected gene expression values. **C, D.** Heatmaps corresponding to upregulated genes in Endothelial-2 cells and fibroblasts comparing the cell location in tumor-context to tumor-adjacent context. The x-axis denotes the genes and y-axis denotes the gene-ontology pathways. The color intensity signifies the fold-change of the DE gene.

**Figure S5. Prostate tumors are enriched in immunosuppressive myeloid cells. A**. Heatmap shows an overview of genes (rows) differentially expressed between myeloid subpopulations of different patients. **B**. Heatmap showing the gene expression dynamics with pseudo-time moving from TIMo to TIMΦ. Representative genes are shown for each cellular state along the cell differentiation (right). Trajectory analysis demonstrates S100A9 (top) and C1QA (bottom) genes expression across pseudotime (left). **C.** Dotplot showing the average expression of the indicated chemokines in the epithelial and stroma subpopulations in our dataset. The color represents scaled average expression of select marker genes in each epithelial subpopulation, and the size indicates the proportion of cells expressing select marker genes. **D**. Enriched Gene Ontology BP categories for the top 200 upregulated genes in each macrophage subpopulations compared to all macrophages. **E**. UMAP embedding for myeloid subpopulations showing the expression of indicated genes. **F**. Detailed annotation of myeloid dendritic cells (mDCs) (left) with marker genes expression shown on UMAP embedding (middle). Right: Stacked bar plot represents the cell proportion of mDCs subpopulations across the three sample fractions.

**Figure S6. Prostate cancer is characterized by T-cell exhaustion and immunosuppressive Treg activity. A**. Boxplot showing the average expression of cytotoxicity score in CD8+ effector T cell subpopulation across the five different samples (Healthy, N-LG, N-HG, T-LG, T-HG). **B**. Boxplot comparing the exhaustion score in CTL-1 (left) and CD8+ effector T cell (right) subpopulations across five different samples. **C**. Boxplot comparing the relative abundance of different lymphoid subpopulations across healthy, adj-normal and tumor fractions. **D**. Boxplot comparing the relative abundance of different lymphoid subpopulations across five different samples. Boxplots in **(A-D)** include centerline, median; box limits, upper and lower quartiles; and whiskers are highest and lowest values no greater than 1.5x interquartile range. Statistical significance was accessed using Wilcoxon rank sum test (*p<0.05, **p<0.01, ***p<0.001). **E**. Heatmap shows average expression of “Treg activity gene signature” (row) in Treg subpopulation in the three different samples (column).

**Figure S7. The prostate cancer TME is enriched ins characterized by exhausted CD56DIM NK cells and activated B cells. A**. Boxplot comparing the relative abundance of NK cells across the 5 different samples. **B**. FGFBP2, GNLY, GZMB and GZMH expression in NK cell subpopulations. **C**. Boxplot comparing the average expression of the exhaustion score between the different NK subpopulations. Statistics significances are accessed using Wilcoxon rank sum test (*p<0.05, **p<0.01). **D**. Boxplot comparing the relative abundance of B cells across the 5 different samples. **E**. Spatial presentation at a high-resolution level using Slide-seqV2 for immune populations in healthy, adj-normal and two tumor tissues collected from a low-grade (Tumor-LG) and high-grade (Tumor-HG) patients. **F**. Dot plot representing key-marker gene expression in immune subpopulations in Slide-seqV2 data. The color represents scaled average expression of marker genes in each cell type, and the size indicates the proportion of cells expressing marker genes. **G**. Stacked bar plot represents the changes in the cell proportion of immune populations obtained from Slide-seqV2 data across the healthy, adj-normal and tumor samples collected from low and high-risk patients.

## Footnotes

1 accession number will available after publication of the manuscript.

