## Supplementary figures and images for "Integrated single-cell and spatial transcriptomic analyses unravel the heterogeneity of the prostate tumor microenvironment"

### Supplementary Figures

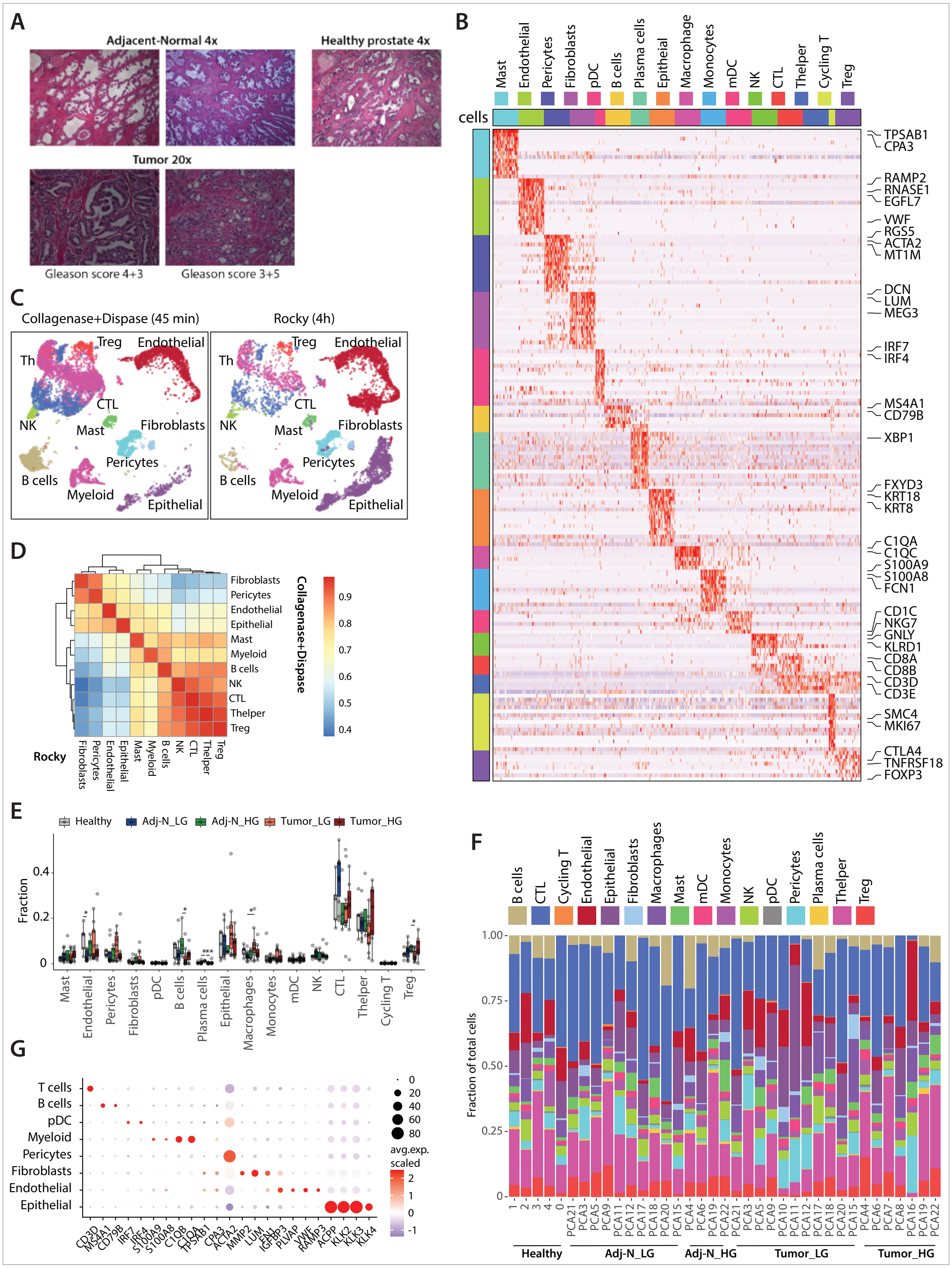

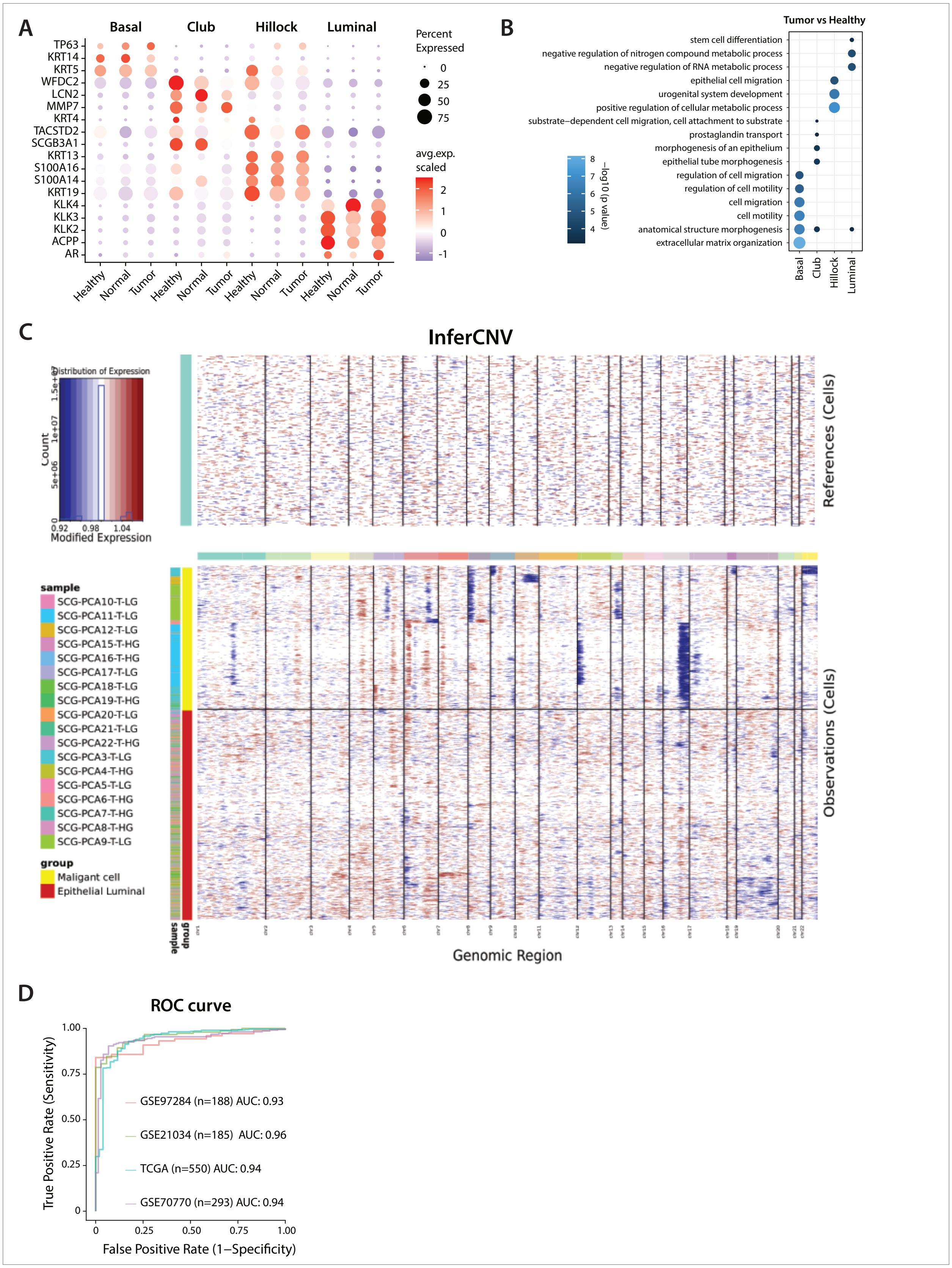

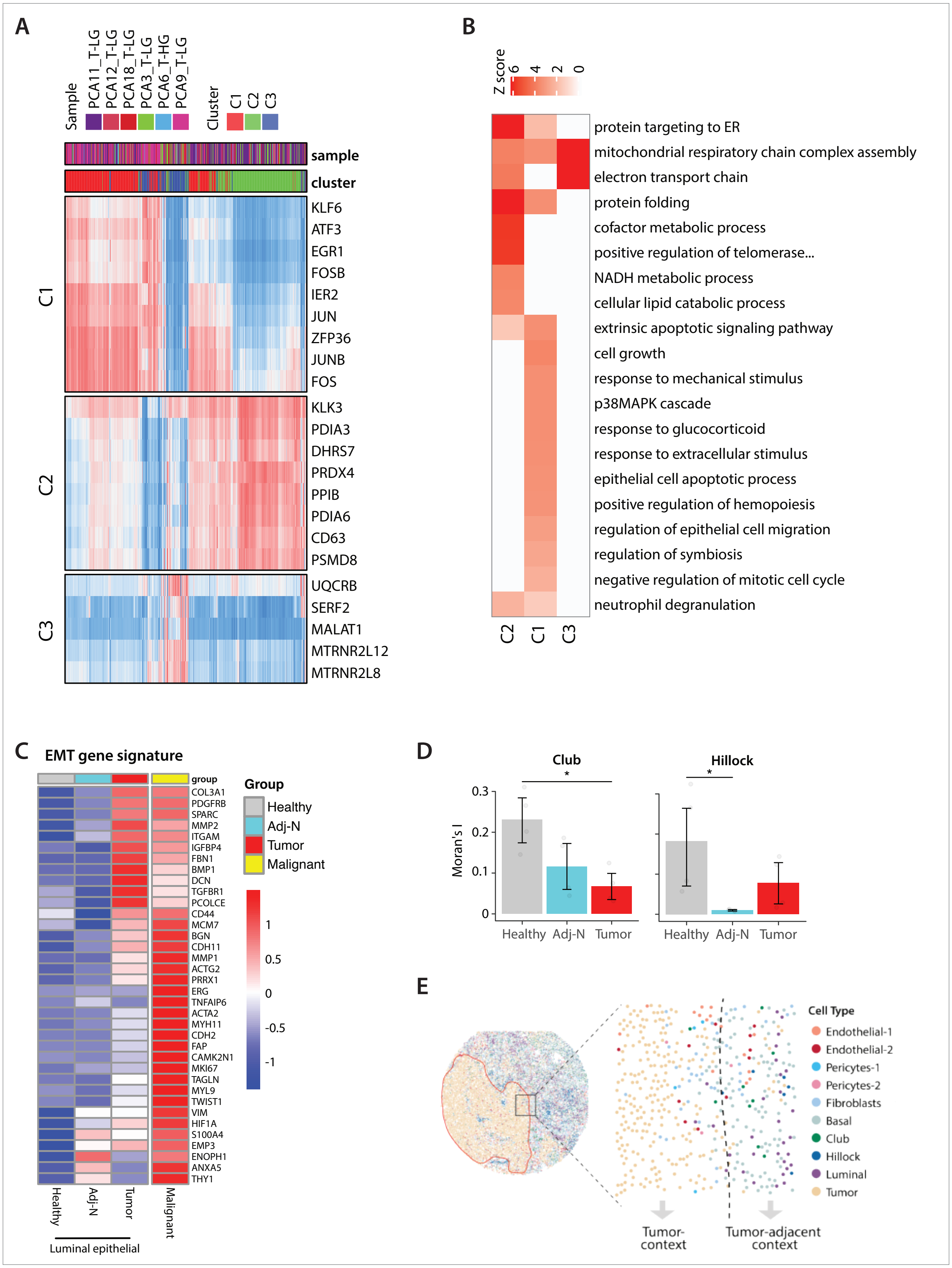

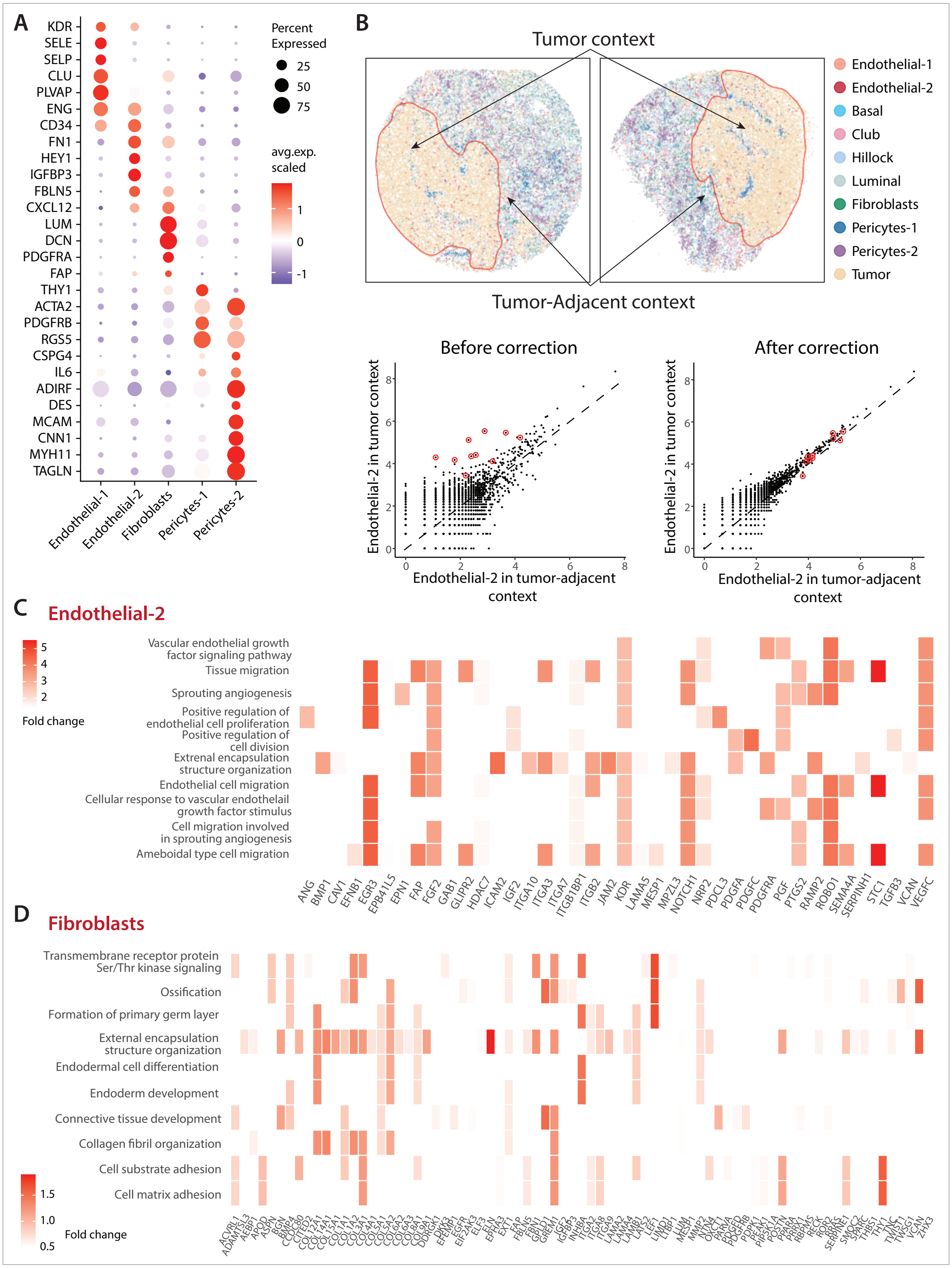

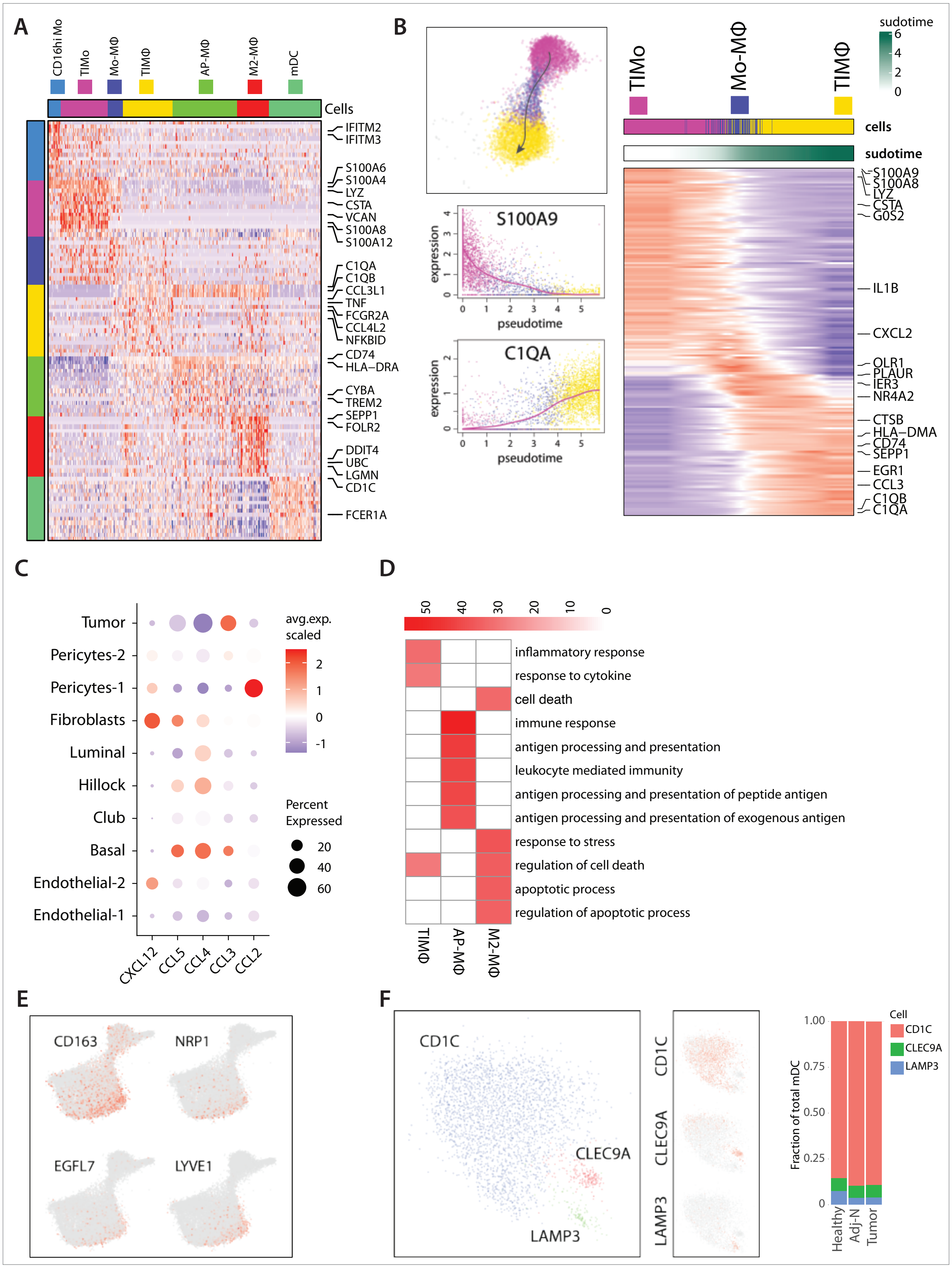

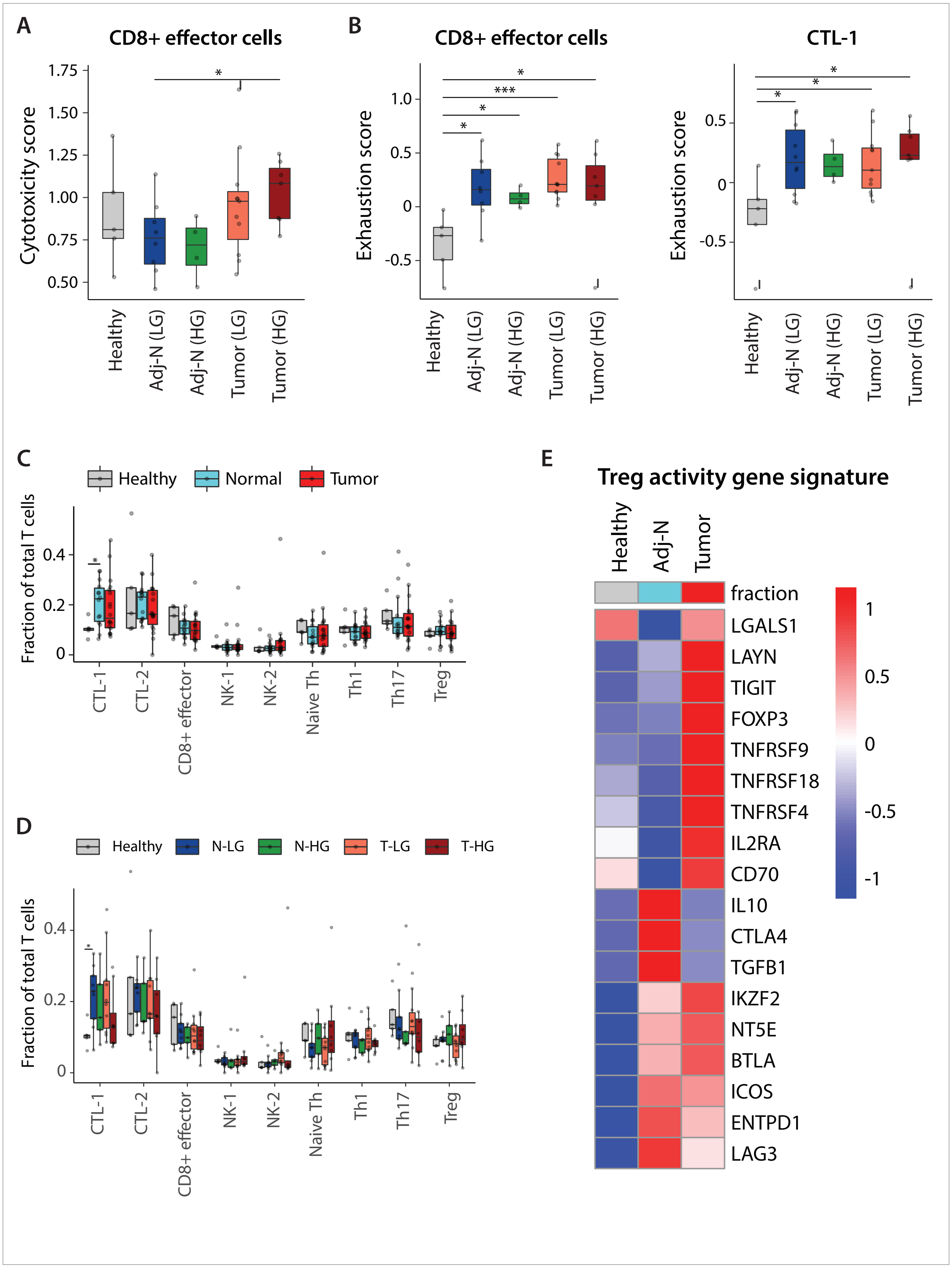

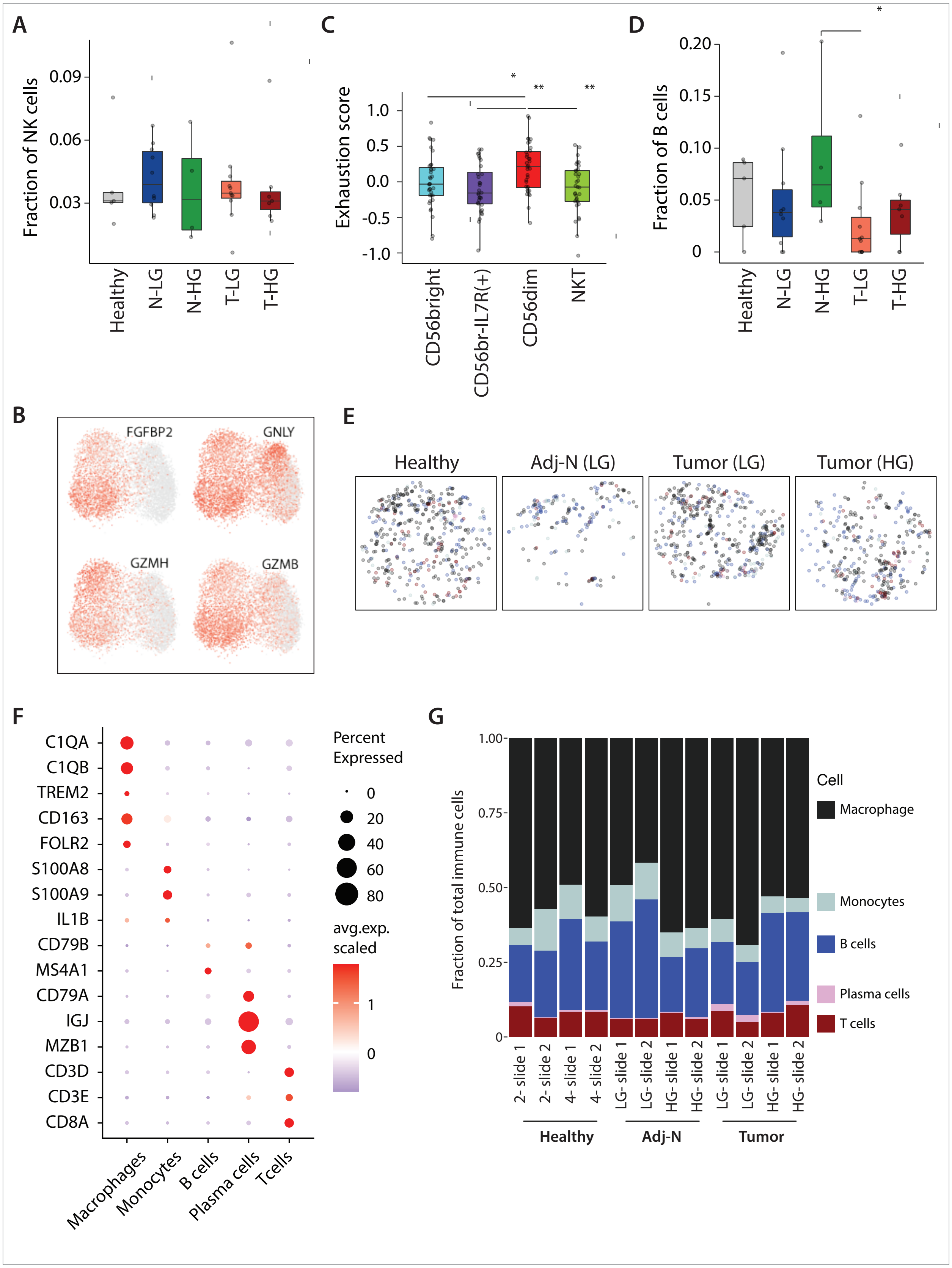
