## Supplementary Note for "Integrated single-cell and spatial transcriptomic analyses unravel the heterogeneity of the prostate tumor microenvironment"

### Supplementary Note: Context-dependent differential expression with linear admixture correction

**Hirz & Mei *et al.***

#### Abstract

Slide-Seq [1] data captures transcriptional profiles of tissue sections in a spatially-resolved manner. A key advantage of such approach is the ability to examine how the expression state of different cell types depends on their context. For instance, the state of an immune cell may differ depending on whether it is found in local inflammatory or non-inflammatory environment. Here we describe an approach for carrying out such tests based on Slide-seqV2 platform. The code repository is available in https://github.com/kharchenkolab/slideseqde

#### Introduction

Slide-seqV2 relies on mRNA hybridization to barcoded bead arrays. The beads are packed on a surface of a glass slide in a circular pattern, forming a "puck". The beads are approximately $10\mu m$ in size and are packed so densely that the distance between bead centers is comparable. Despite such high spatial resolution of the features, inferring transcriptional state of individual cells is challenging due to the fact that multiple cells may contribute to an individual barcoded bead. Figure 1 demonstrates the challenge. A cell may physically overlap more than one bead, and lateral mRNA diffusion during hybridization can further spread mRNA from a given cell to nearby beads. As a result, transcriptional profiles assessed on an individual bead are likely to report a mixture of material from different cells.

A number of computational methods have been developed recently to identify "pure" beads - those with material coming predominantly from a single cell type[2,3]. Such "deconvolution" methods can provide good certainty in the identity of the dominant cell type, however, cannot necessarily discern the detailed transcriptional state of a cell from admixed profiles. Such detailed transcriptional features are central to analysis of how the state of a cell is influenced by its tissue context. For example, consider the state of fibroblast cells in two different contexts (Figure 1a): Tumor context, dominated in its composition by the presence of tumor cells, and Tumor-adjacent context which in addition to fibroblasts contains epithelial and endothelial cells. Even for the beads predominantly capturing material from fibroblast cells (red bead, 1b), the transcriptional profiles captured by the beads will systematically differ between the two context, with the fibroblast beads in the Tumor context capturing substantial signal from adjacent tumor cells, while firboblast beads in the Tumor-adjacent context will capture admixture from epithelial and endothelial neighbors.

To accurately evaluate the transcriptional difference of a given cell type between two contexts, one must correct for the systematic differences in the admixtures. To do so, we use a linear mixture model to estimate coefficients of composition of different cell-types present in the two contexts and correct for them. To simplify the computational problem, we assume that the mixture from the cells to their nearby bead happens in a linear manner, therefore the measured gene expression of each bead is a linear mixture of the nearby cells. Instead of approaching the problem at a single-bead level, we consider average profiles of all beads of a given type in a given context (e.g. by forming pseudo-bulk profiles for all "fibroblast" beads in Tumor context). This is sufficient for answering questions about average differential expression between contexts, and carries two notable advantages. First, pseudo-bulk formulation reduces uncertainty and computational burden. More importantly, together with a linear admixture assumption mentioned above, such formulation allows to avoid the issue of estimating pure admixture profiles. For instance, if endothelial cells exist in both contexts, they may contribute to the target firboblast beads in both contexts and hence would need to be taken into account when correcting the difference. However, the true expression profile of the endothelial cells is also unknown and challenging to estimate, as even "pure" endothelial beads will carry admixture from other cell types, including for example, tumor cells. This sets up a circular dependency that would be difficult to resolve. However, as we will show, under a linear admixture assumption, all such secondary admixtures will cancel out.

#### Computational Pipeline

##### Annotation and Segmentation

The initial annotation of the Slide-seq beads is carried out by an existing tool RCTD[2], based on annotated scRNA-seq data. Given an annotated spatial dataset, a spatial "context" is defined depending on the organization of the tissue and the biological question at hand. In the current manuscript we are interested in investigating the effect of Tumor micro-environment (TME) on

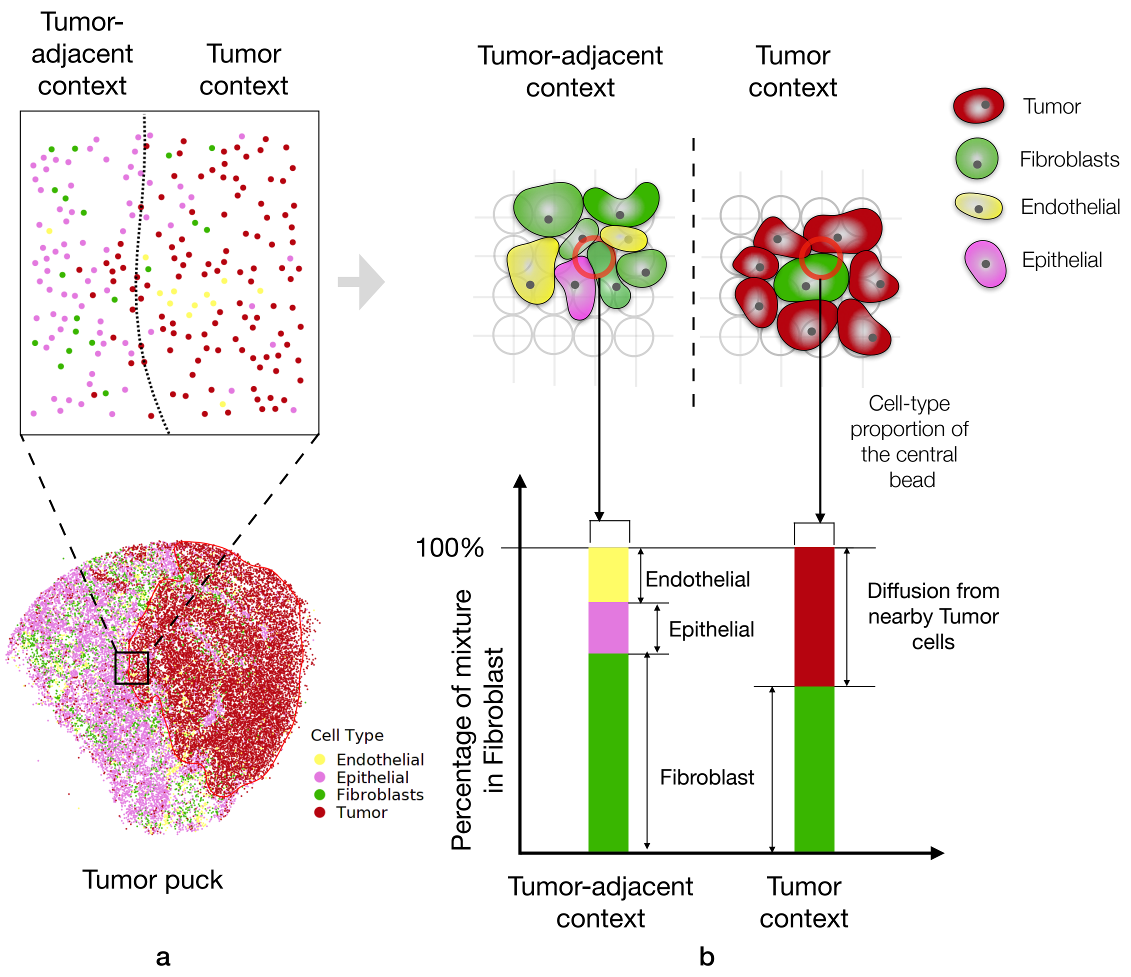


**Figure 1.** Segmenting target cell-type (Tumor) by fitting a kernel density estimator around the spatial locations of the target cell type.

other cell-types. We, therefore, defined the contexts on the basis of the localization of the Tumor cells. As Figure 2 shows, the segmented region (within red boundary) is defined to be the "tumor context", where as the region outside that boundary is defined as as "tumor-adjacent context". Please note that depending on the biological question one sets out to ask, the definition of context and the exact way of determining the segments will change. Contexts may also be spread out, based on local neighborhood properties instead of large-scale segments.

In order to set up the context-dependent differential expression problem, we require three inputs:

1. Gene expression profiles of different cells, formally defined as a matrix $M$ of integer values, of dimension $m \times n$ where there are $m$ cells and $n$ genes
2. Corresponding cell-type annotation for each cell denoted by a set $\mathscr{C}$
3. A set of contexts $D$, defined as a map from the cell id to a discrete variable denoting the context. For example, Figure 2a shows the annotated Slide-seqV2 experiment from the main manuscript, consisting of 10 cell-types, and 2 contexts, namely tumor-adjacent context and tumor context.

Given such a formalization, we first create a composite class for each cell, combining the cell-type of a cell and the context assigned to the cell. As each cell-type can be potentially present in any context, the combined classification will create $|\mathscr{C}||D|$ categories. Next we create pseudo-bulk profiles by summing up (for each gene) the molecules detected in all of the cells within one of the $|\mathscr{C}||D|$ categories. This operation would produce the matrix $M'$ of dimension $n \times |\mathscr{C}||D|$. Note that each column vector of this matrix denotes the pseudo-bulk expression of the cells for a particular cell-type under a specific context. As the column vector of this matrix is of special interest to us, we define the column vector of $M'$ corresponding to cell-type $c_i$ and context $d_j$ as $\kappa_{c_i,d_j}$.

Given a set of contexts and a particular "target" cell-type, we set up a linear model specific to the "target" cell-type. Using the same notations, given 2 contexts $\{d_1, d_2\}$, we first consider a subset of the columns of $M'$, to only include the cell-types within these contexts. Let's denote the reduced matrix as $M'_{d_1,d_2}$ of dimension $n \times 2|\mathscr{C}|$.

#### Regression based correction

To find out the differentially expressed genes for the target cell-type $c_i$ in context $d_1$, when compared to $d_2$, we calculate the following quantities: a matrix $L$ of dimension $n \times (2|\mathscr{C}| - 1)$ with all the columns of $M'_{d_1,d_2}$ except the one that corresponds to the cell-type $c_i$ and the context $d_1$; a vector $\kappa_{c_i,d_1}$ of length $n$, containing the gene expression for the cell-type $c_i$ in the context $d_1$. Using $L$ and $\kappa_{c_i,d_1}$ we then seek a vector $\eta$ of length $2|\mathscr{C}| - 1$ by solving the following constrained optimization problem:

$$\min_{\eta} \|L\eta - \kappa_{c_i,d_1}\| \quad \text{s.t.} \begin{cases} -\infty \leq \eta_j \leq 0 & \text{if } j \in d_2 \text{ and } j \notin c_i \\ 0 \leq \eta_j \leq \infty & \text{Otherwise} \end{cases} \tag{1}$$

The optimization described in the equation 1 can be solved by a bounded-value least square optimization, which we perform by using the `bvls`[4] package in R. The estimated $\eta$ is then used to correct the pseudo bulk profile of the target cell-type $c_i$, in the contexts $d_1$ and $d_2$ by constructing the regressed profiles $\hat{\kappa}_{c_i,d_1}$ and $\hat{\kappa}_{c_i,d_2}$, respectively. Specifically, we compute two vectors $\eta^+$ and $\eta^-$ of length $|\eta|$ as follows:

$$\eta_j^+ = \begin{cases} \eta_j & \text{if } \eta_j > 0 \\ 0 & \text{otherwise} \end{cases} \tag{2}$$

and

$$\eta_j^- = \begin{cases} \eta_j & \text{if } \eta_j < 0 \\ 0 & \text{otherwise} \end{cases} \tag{3}$$

The $\hat{\kappa}_{c_i,d_1}$ and $\hat{\kappa}_{c_i,d_2}$ are computed

$$\hat{\kappa}_{c_i,d_1} = \kappa_{c_i,d_1} - \eta^- L$$
$$\hat{\kappa}_{c_i,d_2} = \eta^+ L$$

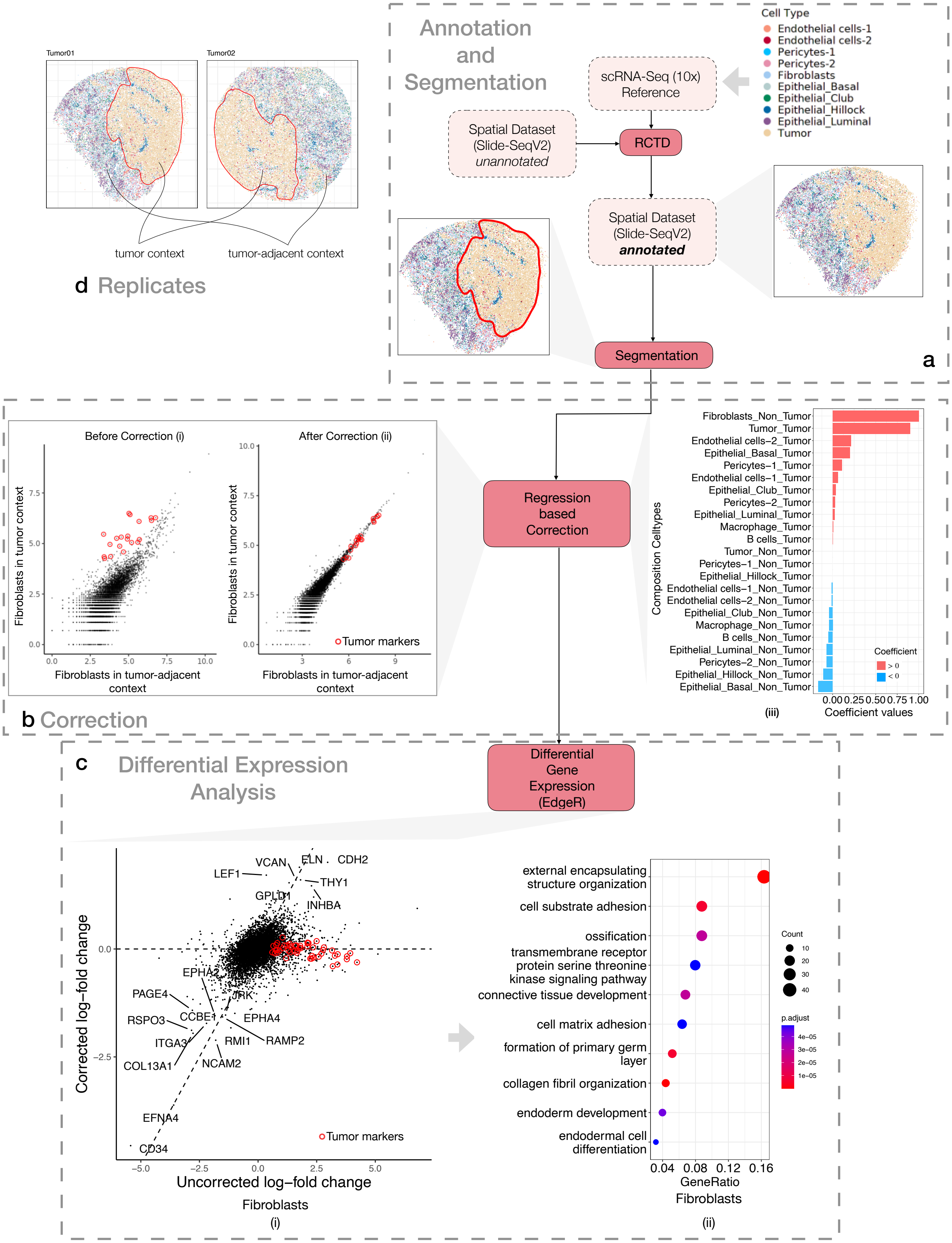


**Figure 2.** The overview of the Slide-seqV2 based computational pipeline.

The plots in Figure 2b show regression-based correction for the prostate cancer datasets from the main manuscript Specifically, we look for expression differences in the fibroblast target cell type, between the tumor context and the tumor-adjacent contexts. The barplot in Figure 2b(iii) shows the coefficients $\eta$ for the cell types in each of the two contexts. Figure 2b shows two scatter plots. Each point corresponds to a gene. The left scatter plot shows expression of each gene in $\kappa_{\text{Fibroblasts,tumor-adjacent}}$ (x-axis) vs $\kappa_{\text{Fibroblasts,tumor}}$ (y-axis). In these uncorrected pseudo-bulk profiles we see inflated expression values for tumor cell markers (denoted by red circles, e.g. TMEFF2, NPY, ERG etc.) demonstrating the phenomenon shown in Figure 1b(i). The other scatter plot ( 1b(ii)) shows analogous contrast between contexts using corrected gene expressions $\hat{\kappa}_{\text{Fibroblasts,tumor-adjacent}}$ vs $\hat{\kappa}_{\text{Fibroblasts,tumor}}$, where the Tumor markers are no longer showing up as being significantly different in their expression.

##### Differential Expression

To test for differentially expressed genes, the corrected profiles $\hat{\kappa}_{c_i,d_1}$ and $\hat{\kappa}_{c_i,d_2}$ are passed to `EdgeR`[5]. When there are more than one Slide-seq replicate present, we apply the regression-based correction for individual replicate before running the DE tool. If run without replicates, the biological variation parameter (`bcv`) within the `EdgeR` is set to 0.1.

Figure 2c shows the results obtained from `EdgeR`[5] based on the corrected Fibroblast profiles from the two HG tumor pucks (Figure 2d). We have also run `EdgeR` with the uncorrected profiles to trace the improvements achieved from the correction. The scatter plot on the left side of Figure 2c(i) shows the log-fold change computed by `EdgeR` with the uncorrected (x-axis) and the corrected profiles (y-axis). Similar to Figure 2b, the tumor cell marker genes are marked by red circles. The log-fold change for the tumor marker genes is reduced with the corrected profiles. Once computed, the top differentially expressed genes could be used for gene enrichment and other downstream analysis (Figure 2c(ii)).
